# Periodontal pathogen-derived extracellular vesicles promote EGFR-dependent malignant traits in human pancreatic cancer cells

**DOI:** 10.64898/2026.01.29.700732

**Authors:** Takehiro Yamaguchi, Ryoma Nakao, Masayuki Shiota, Kimihiro Abe, Yukihiro Akeda

## Abstract

Extracellular vesicles (EVs) produced by periodontopathic bacteria have been implicated in systemic diseases, yet their mechanistic impact on pancreatic cancer (PAC) cell behavior remains poorly defined. Here, we show that EVs derived from *Aggregatibacter actinomycetemcomitans* (Aa), but not from *Porphyromonas gingivalis* or *Fusobacterium nucleatum*, robustly induce epithelial–mesenchymal transition (EMT), migration, and invasion in human PAC cells. These effects require the Aa genotoxin cytolethal distending toxin (Cdt), which is selectively packaged into Aa-derived EVs and efficiently delivered to host cells. EV-associated Cdt induces DNA damage, triggers a senescence-like transcriptional program, increases EGF sensitivity, and activates the EGFR–ERK/Akt signaling axis independently of TGFβ signaling, thereby promoting malignant traits in vitro. Notably, these responses occur in human but not murine PAC cells, revealing species-specific susceptibility to bacterial genotoxins. Together, these findings reveal a previously unrecognized cross-kingdom mechanism by which periodontopathic bacterial EVs promote metastatic traits in human pancreatic cancer cells through Cdt-mediated DNA damage and EGFR–ERK/Akt signaling.

## Introduction

Pancreatic cancer (PAC) is among the top ten causes of cancer-related death (*1*, *2*). Owing to limited advances in therapeutic strategies and the challenges associated with early detection, PAC has a dismal prognosis, with a 5-year survival rate of approximately 10% (*3*, *4*). A hallmark of PAC is its highly invasive nature, which is primarily driven by epithelial–mesenchymal transition (EMT) (*5*). EMT represents a key process in malignant transformation, endowing cancer cells with migratory and invasive capacities that promote local tumor progression and systemic metastasis (*6*).

In addition, PAC is characterized by a unique tumor microenvironment (TME) composed of dense stromal tissue, poor vascularity, and immunosuppressive cell populations, all of which contribute to therapeutic resistance and aggressive disease progression (*3*, *7*). The combined effects of EMT induction and the desmoplastic TME often lead to diagnosis at advanced, inoperable stages, further underpinning the poor prognosis (*5*, *8*).

While established risk factors for PAC include smoking, obesity, excessive alcohol consumption, and a family history of the disease, periodontal disease has recently emerged as a novel and potentially modifiable risk factor for PAC (*3*, *4*). Periodontal disease, driven by infection with periodontopathic bacteria, affects more than one billion people globally and remains a major cause of tooth loss (*9*, *10*); it is also linked to several noncommunicable diseases (NCDs), such as cardiovascular disease, diabetes, and certain cancers (*11*). Recent studies suggest an association between periodontal disease and the development of PAC. Among the diverse array of periodontopathic bacteria, *Porphyromonas gingivalis* (Pg), *Fusobacterium nucleatum* (Fn), and *Aggregatibacter actinomycetemcomitans* (Aa) are particularly associated with PAC (*12*, *13*). However, the mechanisms by which these oral pathogens—originating far from the pancreas—contribute to pancreatic cancer, from initiation to progression, remain largely unknown.

Nearly all bacteria release nanoscale extracellular vesicles (EVs), which are structurally and functionally analogous to eukaryotic exosomes (*14*, *15*). These bacterial EVs carry proteins, nucleic acids, and lipids and act as vehicles for the delivery of virulence factors and toxins (*16*). Exosome-mediated intracellular and interorgan communication has been increasingly recognized as a key mechanism in the pathogenesis of diverse NCDs, including cancer and cardiovascular disorders (*17*, *18*). Analogously, periodontopathic bacteria-derived EVs (perio-EVs) may play a central role in mediating host–pathogen interactions and in the development of periodontal disease-associated NCDs (*19–21*). For instance, Fn-derived EVs (Fn-EVs) promote bacterial adhesion to colorectal epithelial cells and facilitate colorectal cancer development (*22*). Despite these findings, no evidence has yet demonstrated a role for perio-EVs in PAC progression.

In this study, we focused on the potential contribution of perio-EVs to the progression of PAC rather than its initiation. Among EVs from the PAC-associated bacteria Pg, Fn, and Aa, we found that Aa-derived EVs (Aa-EVs) specifically promote the metastatic phenotypes of human PAC cells by promoting cell migration and invasion in a species-dependent manner. Furthermore, we identified cytolethal distending toxin (Cdt), a DNase-active holoenzyme conserved in many gram-negative bacteria, as the critical factor within Aa-EVs responsible for this effect. Cdt in Aa-EVs selectively induced DNA damage in PAC cells, followed by the activation of epidermal growth factor receptor (EGFR) signaling. Our findings suggest that targeting EGFR signaling could present a promising therapeutic strategy for PAC patients infected with Aa and may also provide a mechanistic foundation for understanding how perio-EVs contribute to the pathogenesis of NCDs in distant organs.

## Results

### Aa-EVs promote metastatic phenotypes in PAC cells

EVs were isolated by ultracentrifugation from the culture supernatants of Pg, Fn, and Aa—periodontopathic bacteria previously reported to be associated with PAC. The successful isolation of EVs was confirmed by field-emission scanning electron microscopy (FE-SEM) (Fig. S1A, left). The size distribution of the EVs was confirmed by a nanoflow cytometer, NanoFCM (Fig. S1A, right), and EV-associated protein expression was analyzed by Coomassie Brilliant Blue staining (Fig. S1B). To explore how these perio-EVs affect PAC cell behavior, we first evaluated their effects on key malignant phenotypes, including cell proliferation, migration, and invasion. Aa-EV treatment markedly enhanced the migratory and invasive capacities of human PAC-derived PANC-1 cells (Figs. 1, A and B), whereas it did not significantly affect cell proliferation (Fig. 1C). In contrast, Pg-derived EVs (Pg-EVs) and Fn-EVs did not affect the proliferation or migration of PANC-1 cells (Figs. 1, A and C). Aa-EVs similarly promoted the invasive characteristics of SUIT-2 cells, which are derived from the liver metastasis of a patient with PAC (Fig. 1D). Unlike in PANC-1 cells, Aa-EV treatment significantly suppressed the proliferation of SUIT-2 cells (Fig. 1E). In epithelial cancers, the acquisition of motility is typically linked to EMT.5 Consistently, Aa-EV treatment upregulated the expression of the mesenchymal markers N-cadherin and vimentin but downregulated the expression of the epithelial marker E-cadherin in cells treated with all three types of EVs (Figs. 1, F to I). Together, these results indicate that Aa-EVs specifically promote the acquisition of metastatic phenotypes in PAC cells, likely through the induction of EMT.

**Fig. 1.**
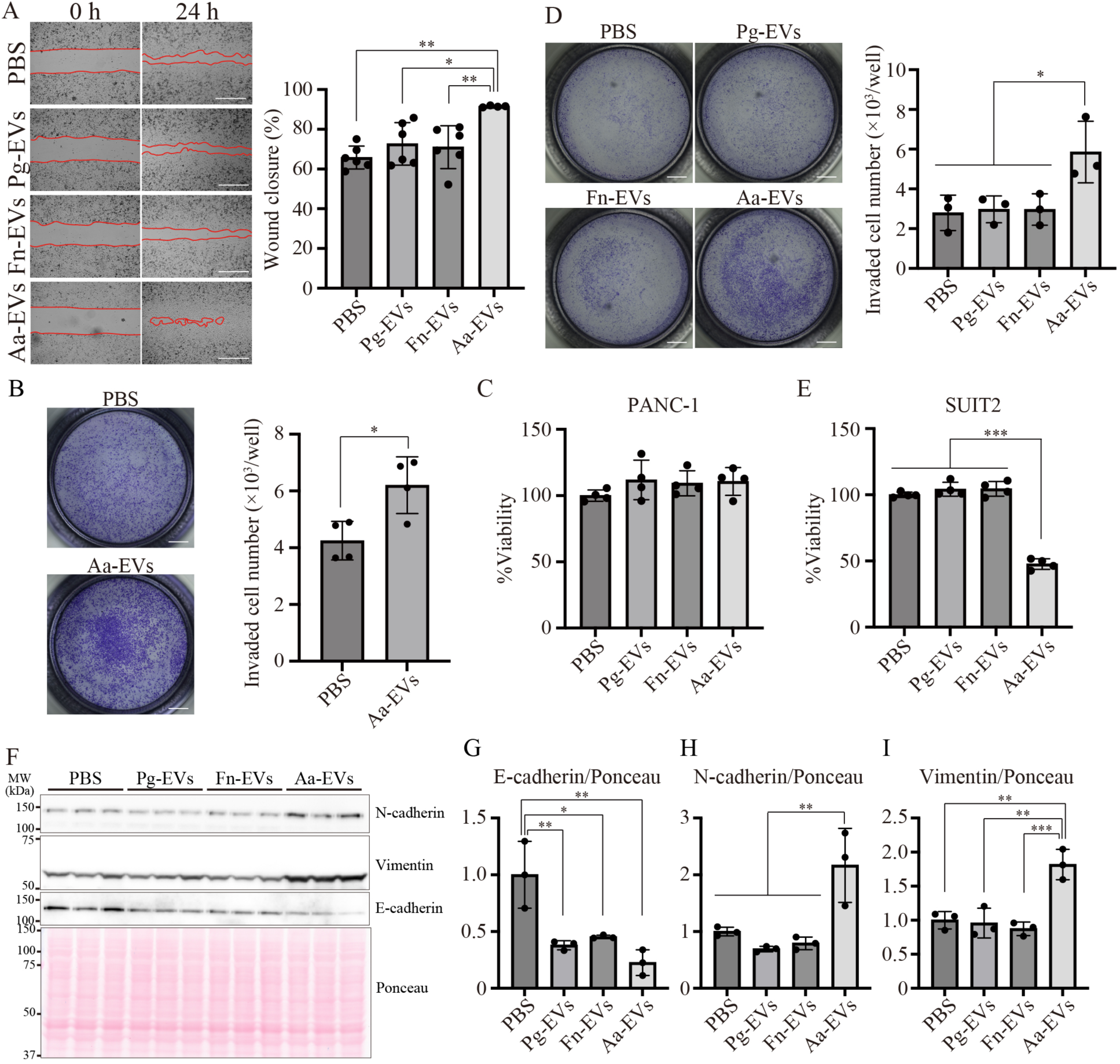
EVs derived from *A. actinomycetemcomitans* (Aa-EVs) promote metastatic phenotypes in human PAC cells through EMT induction. (**A**) Migration assay of PANC-1 cells treated with EVs derived from *Porphyromonas gingivalis* (Pg-EVs), *Fusobacterium nucleatum* (Fn-EVs), or Aa-EVs (left panels) and quantification of the migrated area (right panels; N = 6; three biological and two technical replicates). **(B)** Invasion assay of PANC-1 cells treated with Aa-EVs. Representative microscopy images (left) and quantification (right) are shown. **(C)** The cell viability of EV-treated PANC-1 cells was quantified by a CCK-8 assay (four biological and two technical replicates). **(D)** Invasion assay of SUIT-2 cells treated with Pg-EVs, Fn-EVs, or Aa-EVs (n = 3; three technical replicates). **(E)** The cell viability of EV-treated SUIT-2 cells was quantified by a CCK-8 assay (four biological and two technical replicates). **(F–I)** Western blotting of EMT markers in PANC-1 cells treated with EVs from each bacterium. The quantification of epithelial (G) and mesenchymal (H, I) markers is shown (N = 3; two technical replicates). Representative images from independent experiments are shown. Scale bars: 500 µm (A), 1 mm (B, C). The data are presented as the means ± SDs. *P < 0.05; **P < 0.01; ***P < 0.001 (Tukey’s test). See also Figs. S1 and S2.

### Aa-EVs induce metastatic phenotypes via Cdt but not LtxA

Periodontal disease is observed in a limited range of mammals, including humans, monkeys, dogs, and cats, but not in rodents. Some periodontopathic bacteria are shared across species, such as *Treponema denticola*, Fn, and *Tannerella forsythia* (*23–25*). In contrast, Aa is host-specific and is found primarily in humans and nonhuman primates (*26*). We hypothesized that differences in host susceptibility to Aa may influence the cellular response to Aa-EVs. To test this hypothesis, we examined whether Aa-EVs could induce metastatic phenotypes in murine PAC-derived KPC cells (*27*). As expected, Aa-EV treatment did not increase the migration of KPC cells (Fig. 2A), despite efficient uptake comparable to that observed in human PAC PANC-1 cells (Fig. 2B). We initially hypothesized that leukotoxin A (LtxA), a well-known Aa virulence factor with strict primate specificity due to its targeting of lymphocyte function-associated antigen 1 (LFA-1, integrin αLβ2), might contribute to the species-specific induction of metastatic phenotypes (*28*). However, EVs derived from a *ltxA*-insertion mutant strain of Aa, in which the expression of LtxA was abolished (Figs. S3, A and B) (*29*), promoted migration to a similar extent as that of wild-type Aa-EVs (Fig. 2C), suggesting that LtxA is dispensable in this context.

**Fig. 2.**
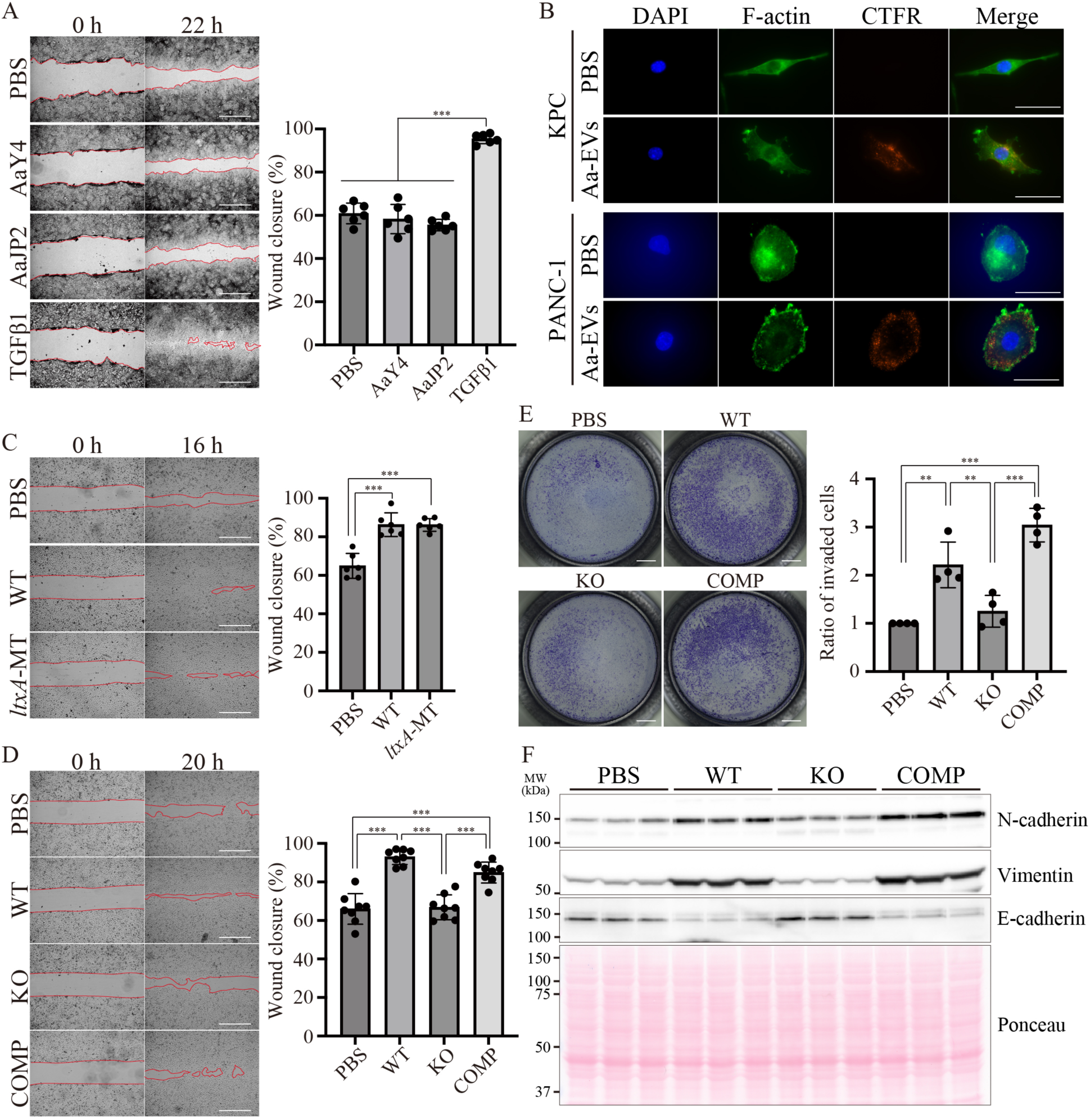
Species-specific induction of metastatic phenotypes by Aa-EVs depends on the cytolethal distending toxin (Cdt). **(A)** Wound-healing assay of murine KPC cells treated with Aa-EVs or TGF-β1. Representative images (left) and quantification (right) (N = 6, two biological and three technical replicates). **(B)** Uptake of CellTrace Far Red (CTFR)-labeled Aa-EVs (red) by PANC-1 and KPC cells. F-actin and nuclei were counterstained with phalloidin (green) and DAPI (blue), respectively. **(C)** Wound-healing assay of PANC-1 cells treated with EVs from wild-type (WT) or *ltxA*-mutant Aa (N = 6). **(D, E)** Effects of *cdtABC* deletion or complementation on Aa-EV-induced migration (D) and invasion (E). PANC-1 (D; N = 8) and SUIT-2 (E; N = 3) cells were analyzed. **(F)** Western blotting of EMT markers in PANC-1 cells treated with EVs from WT, KO, or COMP Aa strains (N = 3). Scale bars: 500 µm (A, C, E), 50 µm (B), and 1 mm (D). The data are presented as the means ± SDs. *P < 0.05; **P < 0.01; ***P < 0.001 (Tukey’s test). See also Figs. S3 and S4.

Aa contains another major virulence factor, cytolethal distending toxin (Cdt), which is composed of three subunits (CdtA, CdtB, and CdtC) (*30–32*). We next investigated the role of Cdt, although the species specificity of Cdt has not been well characterized. We constructed a deletion mutant of the *cdtABC* operon (KO) and a complemented mutant (COMP) in the Aa D7SS strain. EVs from each strain were subsequently isolated (Fig. S3C). Despite apparent alterations in the expression of cellular and MV proteins (Fig. S3D), EVs from Cdt-KO mice failed to induce promigratory and invasive phenotypes in PAC cells. These effects were restored in PAC cells treated with EVs from Cdt-COMP (Figs. 2, D and E). Assessment of EMT marker expression revealed that Aa-EVs increased the expression of the mesenchymal markers N-cadherin and vimentin but downregulated the expression of the epithelial marker E-cadherin in a Cdt-dependent manner (Fig. 2F and figs. S4, A to C). Overall, Cdt, but not LtxA, is essential for the acquisition of metastatic phenotypes in human PAC cells mediated by Aa-EVs through the induction of Cdt-dependent EMT.

### Aa-EV-associated Cdt mediates DNA damage and cellular quiescence

Cdt is a DNase-active holoenzyme conserved among various gram-negative bacteria and is known to induce DNA damage followed by cell cycle arrest in host cells (*31*, *33*).

Previous studies have reported that DNA damage and quiescence promote invasion in specific cancer cells (*34*). We hypothesized that Cdt-induced DNA damage may trigger cellular quiescence and subsequently promote the migration and invasion of PAC cells. When human PAC cells were treated with Aa-EVs, DNA damage was induced in a Cdt-dependent manner, as evidenced by γH2AX staining (Figs. 3, A and B). However, Aa-EVs did not induce DNA damage in mouse PAC-derived KPC cells (Figs. 3, C and D). Time-lapse imaging revealed that cell proliferation was suppressed in both PANC-1 and SUIT-2 cells in a Cdt-dependent manner (Figs. 3, E and F). Although there was no significant difference in the cell viability of PANC-1 cells (Fig. 3G), the cell viability of SUIT-2 cells was suppressed in a Cdt-dependent manner. These data suggest that DNA damage, which is induced by Aa-EV-associated Cdt in a species-specific manner, may play a critical role in the acquisition of metastatic phenotypes of PAC.

**Fig. 3.**
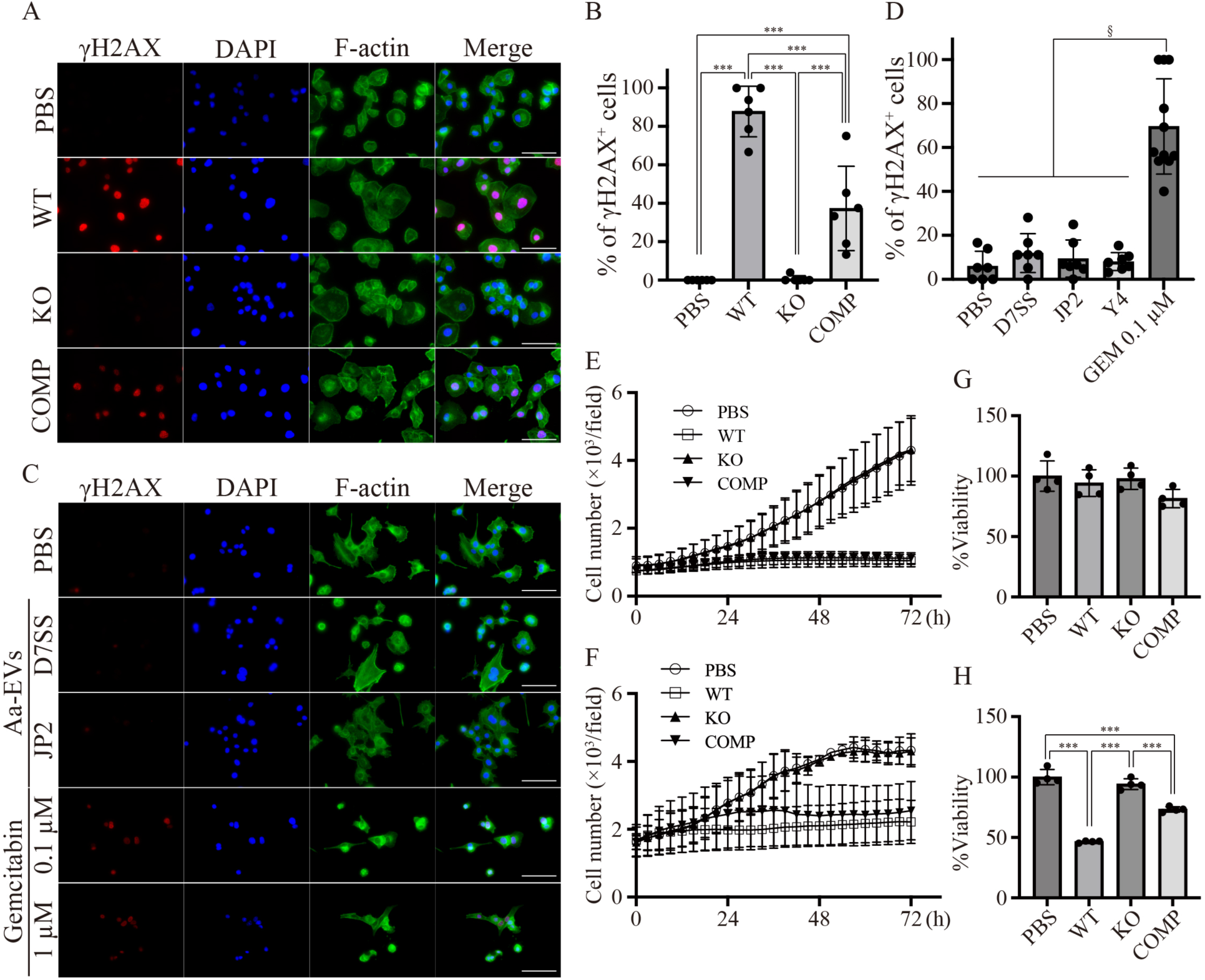
Aa-EV-associated Cdt induces DNA damage and cellular quiescence in a species-specific manner. (A, **C)** Immunofluorescence of γH2AX (red), F-actin (green), and nuclei (blue) in PANC-1 (A) and KPC (C) cells treated with Aa-EVs or gemcitabine. **(B, D)** Quantification of DNA-damaged cells. **(E, F)** Time-lapse proliferation of PANC-1 (E) and SUIT-2 (F) cells treated with EVs from Cdt-mutant Aa strains (N = 6). **(G, H)** Viability assays of PANC-1 (G) and SUIT-2 (H) cells (N = 4). Representative results and images from replicate experiments are shown. Scale bars: 100 µm. The data are presented as the means ± SDs. *P < 0.05; **P < 0.01; ***P < 0.001 (Tukey’s test).

### EV encapsulation of Cdt efficiently induces the metastatic phenotypes

EVs are considered vehicles that effectively transfer cargoes intercellularly between bacteria and bacteria and between bacteria and the host. EVs also protect their cargoes against proteases and nucleases. We hypothesized that, when delivered via EVs, Cdt also effectively induces the metastatic phenotype of PAC; to demonstrate this, we constructed recombinant Cdt (rCdt), which consisted of His6-tagged CdtA, CdtB, and CdtC (Figs. S5, A to D). First, we evaluated the Cdt content of Aa-EVs derived from the His6-tagged Cdt-expressing strain (COMP) using purified recombinant CdtB protein (Fig. S5E).

Quantitative Western blotting revealed that CdtB was present in COMP-derived EVs at 23±16.4 fmol/µg (Fig. 4A). Because COMP-derived EVs promoted the metastatic features of human PAC cells at a concentration of 1 µg/mL (Figs. 2, D and E), the expected effective concentration of EV-associated Cdt was 23.4±16.4 pM. How much non-EV-associated rCdt is required to induce the metastatic phenotypes of PAC cells?

**Fig. 4.**
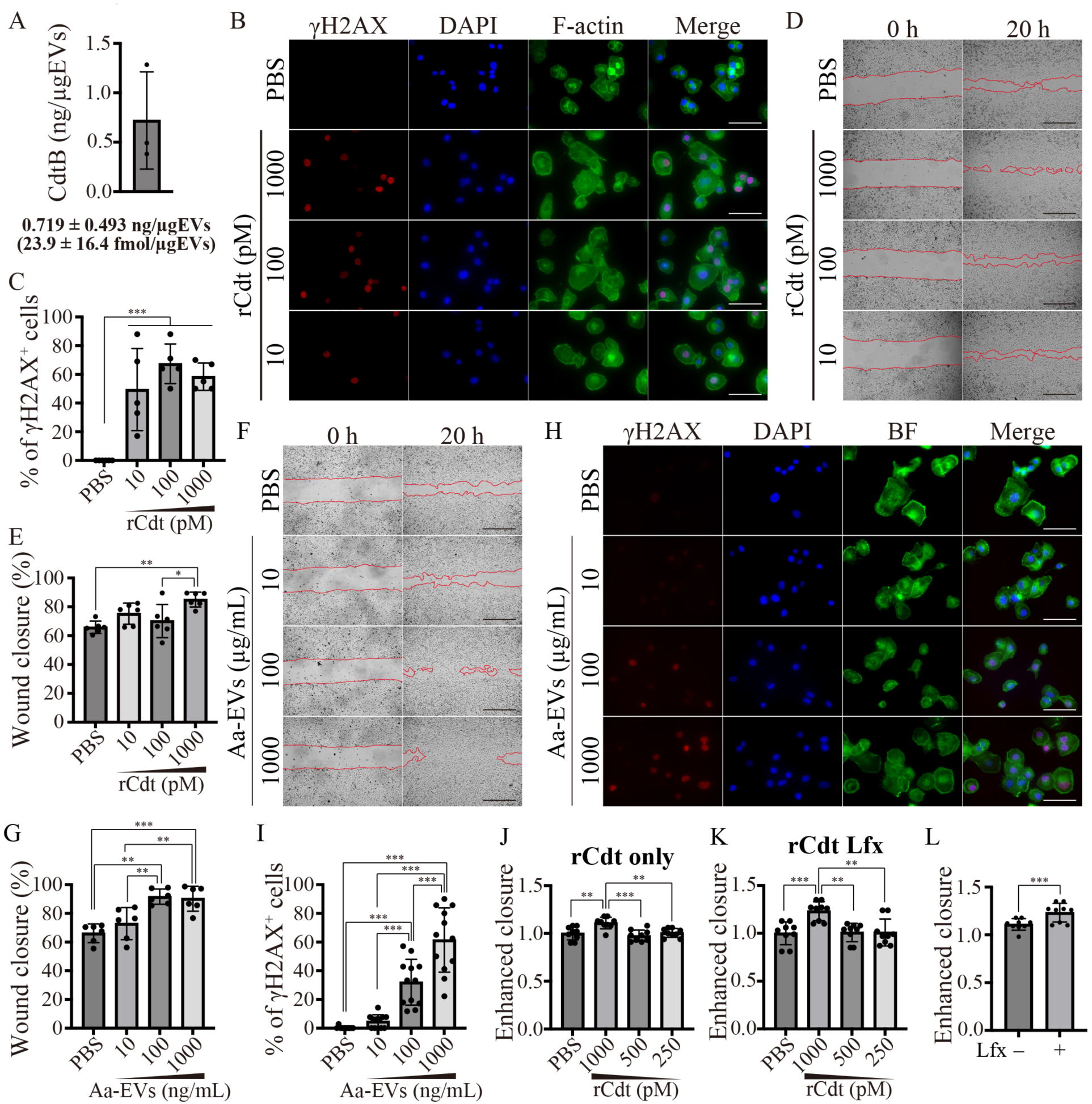
Recombinant Cdt induces DNA damage but does not promote migration in PAC cells. **(A)** Quantification of CdtB content in His₆-tagged Cdt-expressing Aa-EVs. **(B, C)** Concentration-dependent effects of recombinant Cdt (rCdt) on migration in PANC-1 cells. **(D, E)** rCdt-induced DNA damage visualized by γH2AX staining (D) and quantified (E). **(F–H)** Effect of protein lipofection on rCdt-induced migration. Wound-healing assays of rCdt only (F) or rCdt + lipofection (G), with quantification (H). **(I–L)** Concentration-dependent effects of Aa-EVs on migration (I, J) and DNA damage (K, L) in PANC-1 cells. Scale bars: 500 µm (B, I), 100 µm (D, K). The data are presented as the means ± SDs. *P < 0.05; **P < 0.01; ***P < 0.001 (Tukey’s test). See also Figs. S5–S7.

Unexpectedly, rCdt induced DNA damage in PANC-1 cells at concentrations above 10 pM (Figs. 4B and C), similar to the effective concentration of EV-associated Cdt, although rCdtA, rCdtB, and rCdtC could not induce DNA damage even at 100 nM (Fig. S6).

However, the effective rCdt concentration for promoting cellular migration was 1,000 pM (Figs. 4D and 4E), which was relatively higher than that for inducing DNA damage. At concentrations of 100–1,000 µg/mL, Aa-EVs increased the migration of PANC-1 cells (Figs. 4, F and G). In parallel, Aa-EV-associated Cdt can induce DNA damage at concentrations of 100–1,000 µg/mL (Figs. 4, H and I). Aa-EV-associated Cdt can both increase cellular migration and induce DNA damage. Finally, we investigated whether the delivery of rCdt by lipofection could increase cellular migration more effectively. After confirming the successful procedure of lipofection using R-phycoerythrin (Fig. S7A), we determined the effective concentrations of rCdt for increasing cellular migration with or without a lipofection reagent. Both rCdt alone and rCdt with lipofection reagent increased cellular migration at a concentration of 1,000 pM (Figs. 4, J and 4K, and figs. S7, B and C). Lipofection increased the migration of PANC-1 cells treated with rCdt (Fig. 4L) but did not affect the effective concentration of rCdt. These findings suggest that Cdt-induced DNA damage triggers but does not adequately enhance the metastatic phenotypes of PAC cells. These results suggest that Aa-EVs contain unknown enhancers, which may cooperate with Cdt and effectively induce metastatic phenotypes in PAC cells.

### Aa-EVs induce EMT and genomic instability

Next, we sought to clarify why the DNA damage induced by Cdt triggers EMT and promotes the acquisition of metastatic features in PAC cells. To explore the underlying mechanism, we analyzed the gene expression profiles of the Aa-EV-treated PANC-1 cells using RNA-seq (Fig. 5A and fig. S8A). Aa-EV treatment upregulated 887 genes and downregulated 157 genes. Hallmark pathway enrichment analysis of the upregulated genes using gene sets from the Molecular Signatures Database (MSigDB) revealed significant enrichment of gene sets related to DNA damage responses, such as E2F targets, the G2M checkpoint, and the UV response, as well as epithelial–mesenchymal transition (EMT) (Fig. 5B). Among the EMT-related genes, 26—including key regulators of EMT induction, such as *SNAI1* and *THY1*—were upregulated by Aa-EV treatment (Fig. 5C).

**Fig. 5.**
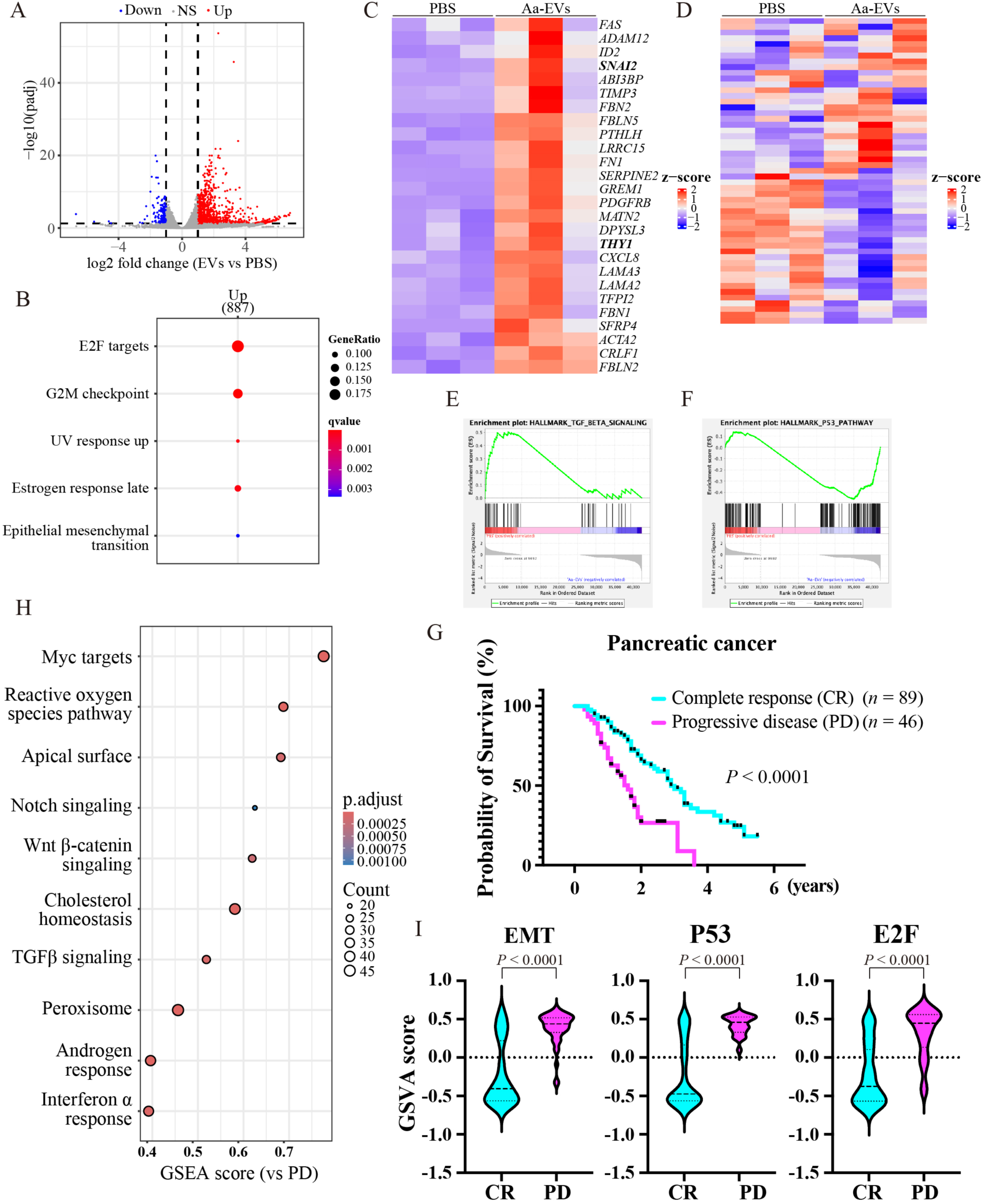
Transcriptomic analysis revealed the activation of DNA damage responses and TGFβ1-independent EMT by Aa-EVs. **(A)** Volcano plot of differentially expressed genes (DEGs) between Aa-EV-treated and untreated PANC-1 cells. **(B)** Hallmark pathway enrichment analysis of upregulated DEGs. **(C, D)** Heatmaps showing changes in EMT-related genes (C) and TGFβ signaling-related genes (D). **(E, F)** GSEA plots of genes associated with TGFβ and p53 signaling pathways. **(G)** Kaplan–Meier survival analysis of pancreatic cancer (PAC) patients with complete response (CR) or progressive disease (PD). **(H)** Dot plot showing gene set enrichment analysis (GSEA) results comparing CR versus PD in PAC patients. **(I)** Violin plots showing gene set variation analysis (GSVA) enrichment scores of EMT-, P53-, and E2F-related pathways between patients who achieved CR and those who achieved PD. See also Figs. S8 and S9.

Although TGFβ is a well-known inducer of EMT (*35*, *36*), treatment with Aa-EV did not activate the TGFβ signaling pathway (Fig. 5D); instead, gene-set enrichment analysis (GSEA) revealed that TGFβ signaling was suppressed (Fig. 5E). In contrast, GSEA revealed the activation of pathways related to DNA damage, including the UV response, the p53 pathway, and apoptosis (Fig. 5F and figs. S8, C to E).

Genomic instability is a hallmark and a driving force of cancer progression. Aa-EV-associated Cdt, a DNase I-like nuclease, induces DNA damage responses, which may in turn promote genomic instability and cancer progression. To assess the clinical relevance of the Aa-EV-induced transcriptomic changes observed in vitro, we analyzed the RNA-seq data of PAC patients available from the GDC Data Portal (https://portal.gdc.cancer.gov). When patients with a complete response (CR) were compared to those with progressive disease (PD), the probability of survival was significantly lower in the PD group (Fig. 5G). GSEA revealed that the TGFβ signaling pathway was activated in the PD group (Fig. 5H). Although GSEA did not detect significant differences in DNA damage-related or EMT pathways, gene set variation analysis (GSVA) revealed strong correlations between PAC malignancy and both EMT induction and activation of DNA damage-related pathways (p53 and E2F) (Fig. 5I), which is consistent with the findings in Aa-EV-treated PANC-1 cells (Figs. 5, B and F, and figs. S8, C and D). Notably, similar activation of DNA damage-related factors, such as E2F targets and G2M checkpoints, was also observed when colorectal cancer patients with CR and those with PD were compared in the GDC Data Portal dataset (Figs. S9, A and B), suggesting that these transcriptomic signatures may represent a common molecular consequence of Cdt exposure across tumor types. Together, these results indicate that Aa-EV-associated Cdt promotes DNA damage-driven transcriptomic reprogramming linked to EMT and cancer progression in PAC cells.

### Aa-EV increases EGF sensitivity and activates EGFR signaling to induce EMT

Our results indicated that Aa-EV treatment induced EMT without activating the canonical TGFβ pathway, which is typically required for the induction of EMT (Figs. 5, D and E, and figs. S8, A to C). To identify alternative pathways responsible for EMT induction, we explored other signaling cascades using an RNA-seq dataset of Aa-EV-treated PANC-1 cells. Previous studies have shown that the DNA damage response is closely linked to epidermal growth factor receptor (EGFR) signaling (*37–39*). EGFR signaling facilitates cell survival against DNA damage (*37*, *38*), and DNA damage itself can increase EGFR signaling (*39*). Our results revealed that Aa-EV treatment increased the phosphorylation of ERK1/2 but not that of Akt—both of which are downstream molecules of EGFR signaling—in a Cdt-dependent manner (Figs. S2, A to C and figs. S4, D to F). Moreover, compared with pretreatment with PBS, pretreatment with Aa-EVs markedly increased the cellular response to EGF stimulation, as shown by the increased phosphorylation of EGFR, ERK1/2, and Akt (Fig.6A). These results suggest that Aa-EV-associated Cdt increases EGF sensitivity and primes EGFR signaling activation in PAC cells.

**Fig. 6.**
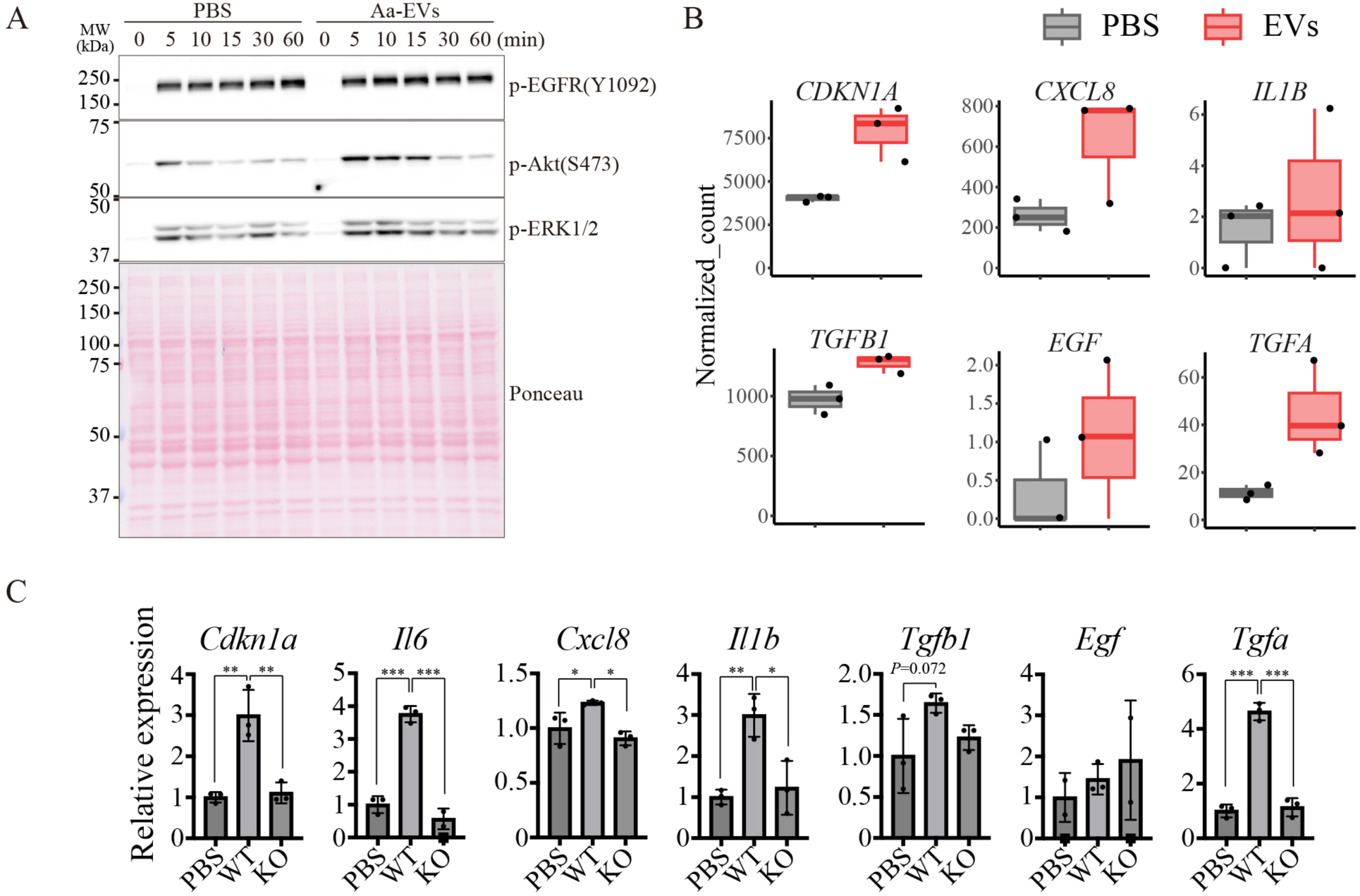
Aa-EVs induce activation of EGFR signaling and a senescence-like transcriptional program. **(A)** Comparison of EGFR signaling activation in EGF-treated PANC-1 cells with or without Aa-EV pretreatment. **(B)** Normalized RNA-seq counts of selected genes (*CDKN1A*, *CXCL8*, *IL1B*, *TGFB1*, *EGF*, and *TGFA*). **(C)** Validation of selected mRNAs by qPCR in Cdt-mutant Aa-EV–treated PANC-1 cells (n = 3). The data are presented as the means ± SDs. *P < 0.05; **P < 0.01; ***P < 0.001 (Tukey’s test). See also Fig. S10.

Evaluation of gene expression using normalized counts from the RNA-seq dataset of Aa-EV-treated PANC-1 cells revealed upregulation of *CDKN1A* and *CXCL8*, which are related to cell cycle arrest and EMT induction, respectively (Fig. 6B). The upregulation of these genes, which was dependent on Cdt, was validated by qPCR (Fig. 6C). The expression of *IL1B* and *IL6*, which are associated with inflammation and cellular senescence, was also upregulated in a Cdt-dependent manner, although the results of the transcriptomic analysis did not reveal these alterations (Figs. 6, B and C). The expression of *TGFB1* was slightly upregulated (less than twofold) in Aa-EV-treated cells, despite the overall suppression of the TGFβ signaling pathway (Figs. 5F and 6B), and qPCR analysis also revealed a trend toward increased *TGFB1* expression (Fig. 6C). Notably, the RNA-seq data revealed that the expression of *TGFA*, a ligand of EGFR, was upregulated by Aa-EV treatment, whereas that of *EGF* alone was not (Fig. 6B). These findings were validated by qPCR, which confirmed that Aa-EV-associated Cdt significantly upregulated *TGFA* expression (Fig. 6C). Consistent with these findings, the phosphorylation of Smad2/3 was not increased by Aa-EV treatment, indicating that the canonical TGFβ pathway remained inactive under these conditions (Fig. S10). Taken together, these findings suggest that Aa-EV-associated Cdt induces DNA damage responses and increases EGF sensitivity, which may lead to activation of the TGFα–EGFR–ERK signaling axis rather than the canonical TGFβ–Smad pathway, thereby promoting EMT and metastatic features in PAC cells.

### Aa-EV-induced migration requires the EGFR–ERK/Akt signaling axis

Finally, we examined whether inhibition of EGFR signaling abrogates EMT induction and increases migratory activity in Aa-EV-treated PANC-1 cells. Treatment with the ERK1/2 inhibitor SCH772984 (SCH, 2 µM) or the Akt inhibitor MK2206 (MK, 2 µM) partially suppressed Aa-EV-induced cellular migration (Figs. 7, A and B). SCH treatment more strongly reduced migration (PBS + SCH vs. Aa-EVs + SCH, 47.77 ± 3.77% vs. 55.16 ± 6.60%) than did MK treatment (PBS + MK vs. Aa-EVs + MK, 42.48 ± 6.07% vs. 58.39 ± 5.17%), which is consistent with the predominant activation of ERK1/2 rather than Akt by Aa-EV treatment (Figs. S2, A to C and figs. S4, D to F). Combined treatment with SCH and MK (2 µM SCH + 2 µM MK), which completely blocked downstream EGFR signaling, abolished Aa-EV-induced migration (Fig. 7C; PBS + SCH + MK vs. Aa-EVs + SCH + MK, 34.36 ± 2.19% vs. 30.83 ± 3.99%, respectively). Moreover, inhibition of EGFR itself by sapitinib (10 µM) completely suppressed the increase in migration induced by Aa-EVs (Fig. 7D; PBS + sapitinib vs. Aa-EVs + sapitinib, 34.31 ± 8.16% vs. 36.62 ± 6.63%, respectively). Together, these findings support a model in which periodontopathic Aa-derived EV-associated Cdt induces DNA damage in pancreatic cancer cells, leading to enhanced EGFR–ERK/Akt signaling and subsequent EMT induction, thereby promoting metastatic traits (Fig. 8).

**Fig. 7.**
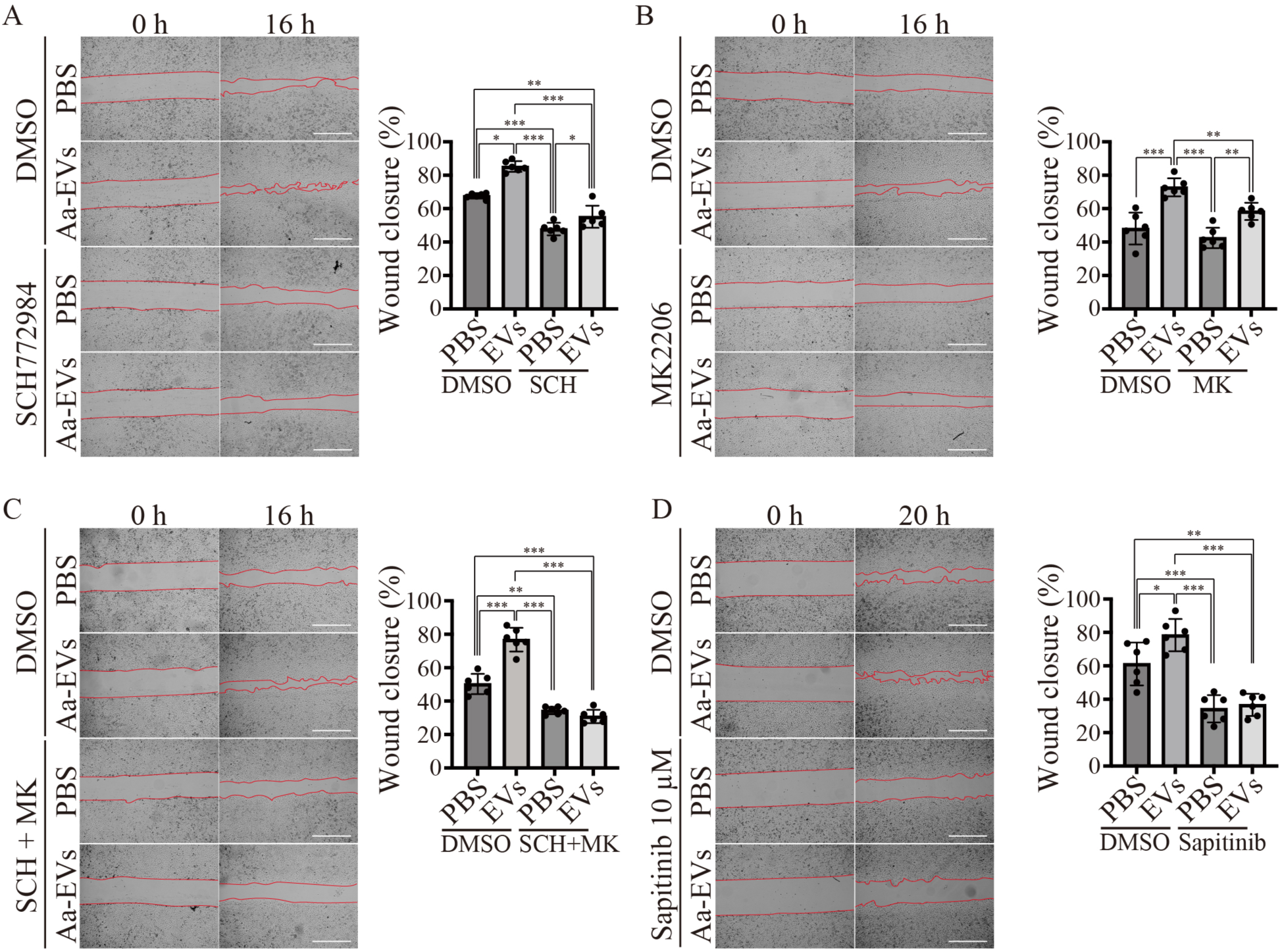
Aa-EV-induced EMT requires EGFR–ERK/Akt signaling. (A–C) Effects of ERK1/2 inhibition (SCH772984; 2 µM), Akt inhibition (MK2206; 2 µM), and dual inhibition (SCH + MK) on Aa-EV-induced migration (n = 6). **(D)** Effect of the pan-EGFR inhibitor sapitinib (10 µM) on Aa-EV-induced migration (n = 6). Representative microscopy images (left panels) and quantification (right panels) are shown. Scale bars: 500 µm. The data are presented as the means ± SDs. *P < 0.05; **P < 0.01; ***P < 0.001 (Tukey’s test).

**Fig. 8.**
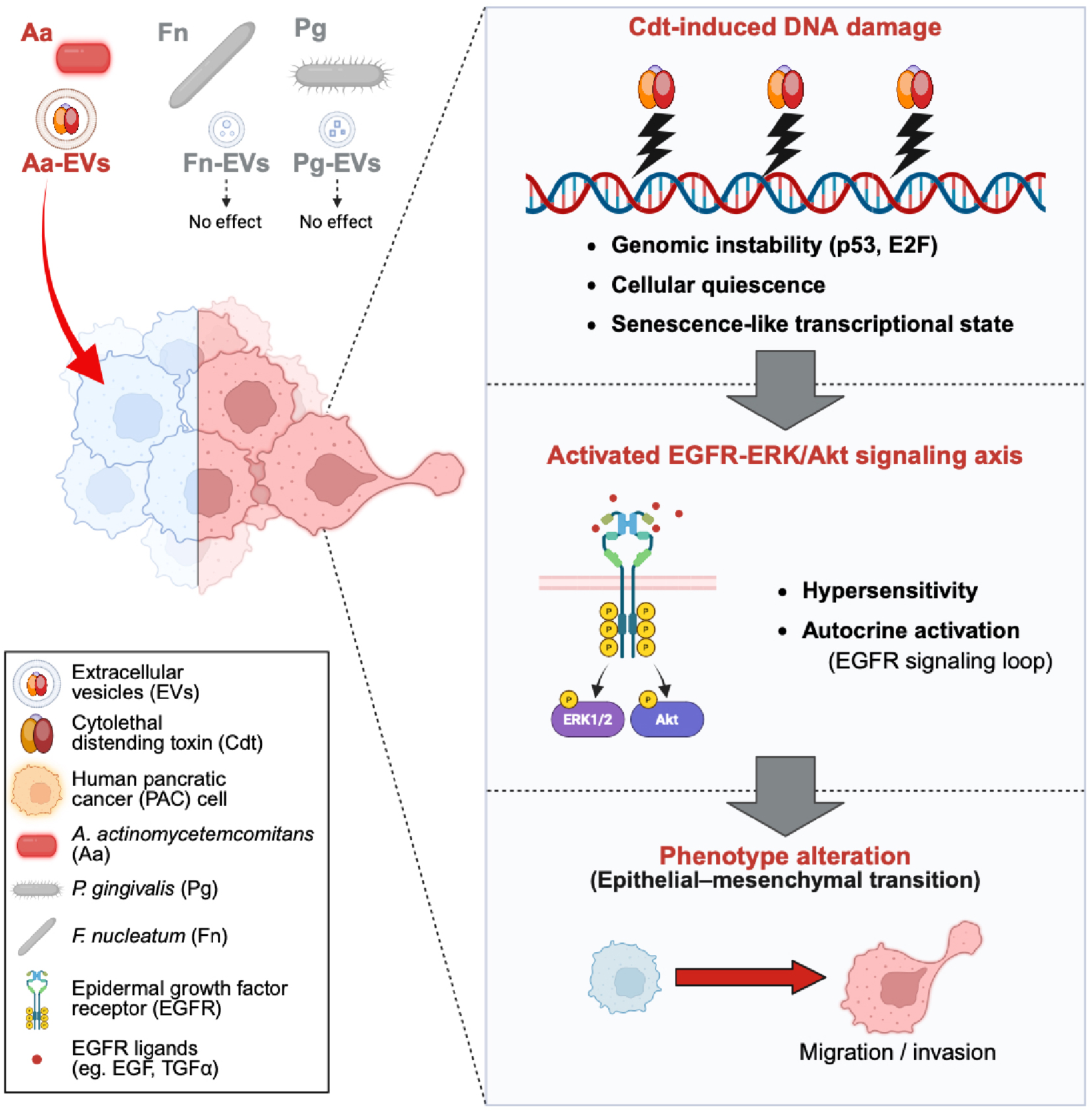
Proposed model of Aa-EV-associated Cdt-mediated promotion of metastatic traits in pancreatic cancer cells. Aa-derived extracellular vesicles deliver cytolethal distending toxin (Cdt) into human pancreatic cancer cells, inducing DNA damage and enhancing EGFR–ERK/Akt signaling in a TGFβ-independent manner. This signaling cascade promotes epithelial–mesenchymal transition (EMT) and the acquisition of metastatic traits.

## Discussion

In this study, we identify a previously unrecognized mechanism by which extracellular vesicles (EVs) derived from the periodontal pathogen *Aggregatibacter actinomycetemcomitans* (Aa) promote metastatic traits in human pancreatic cancer cells. Specifically, we show that Aa-derived EVs deliver the genotoxin cytolethal distending toxin (Cdt), which induces DNA damage-driven transcriptional reprogramming, enhances EGF sensitivity, and activates the EGFR–ERK/Akt signaling axis, thereby promoting epithelial–mesenchymal transition (EMT) and invasive behavior.

Cdt is a DNase I-like enzyme that induces DNA double-strand breaks and subsequent activation of the DNA damage response (DDR) in eukaryotic cells (*30–33*, *40*, *41*). In our study, Aa-EV-associated Cdt triggered γH2AX foci formation and the upregulation of *CDKN1A*, *IL1B*, *IL6*, and *CXCL8*, all of which are canonical hallmarks of cellular senescence (*42*, *43*). Although we did not assess β-galactosidase activity—a classical hallmark of cellular senescence—the transcriptional signature together with Cdt-dependent growth suppression indicated that Aa-EVs induce a senescence-like state in PAC cells. While cellular senescence is typically a tumor-suppressive response to DNA damage, senescent or senescence-like cells often acquire a senescence-associated secretory phenotype (SASP), which can exert protumorigenic effects through chronic secretion of cytokines and growth factors (*44–46*). Therefore, the Cdt-induced senescence-like state may represent a protumorigenic cellular state. Although our present study focused on the promotion of metastatic phenotypes, these findings raise the intriguing possibility that Aa-EVs might also contribute to the initiation of PAC.

We also found that Aa-EV-associated Cdt induced DNA damage and EMT exclusively in human PAC cells but not in murine PAC cells. While the species specificity of another Aa virulence factor, leukotoxin A (LtxA), has been well characterized—it selectively targets primate LFA-1—the species-dependent activity of Cdt has not been reported previously (*47–49*). This study provides the first evidence demonstrating the species-specific action of Cdt. These findings suggest that Cdt requires human- or primate-specific host factors for its cellular uptake or for DNase activation in target cells. The species restriction of Cdt activity also explains the difficulty of recapitulating Aa-EV-mediated oncogenic processes in standard rodent models. Future studies using nonhuman primate models or humanized mice will be essential for evaluating the in vivo relevance of these findings.

Key mechanistic insight from our work is that Aa-EV-associated Cdt induces EMT independent of the canonical TGFβ–Smad pathway, which is typically required for EMT induction (*35*, *36*). Instead, Aa-EV treatment increased EGF sensitivity and activated the EGFR–ERK/Akt signaling axis in a Cdt-dependent manner. Among the EGFR ligands, *TGFA* was significantly upregulated, whereas *EGF* itself was not. This Cdt-induced increase in EGFR signaling may be partly explained by the suppression of *MIG6* (also known as *ERRFI1*), a negative feedback regulator of EGFR whose expression is known to be repressed by DNA damage (*39*). Such downregulation of *MIG6* expression could relieve EGFR inhibition, thereby sensitizing PAC cells to EGF stimulation and sustaining oncogenic signaling. These results suggest that DNA damage caused by Cdt may trigger compensatory activation of EGFR signaling, which is consistent with previous reports that the DDR and EGFR pathways are mutually reinforcing in cancer cells (*37–39*). While

Cdt-induced DNA damage is sufficient to trigger EMT-related transcriptional reprogramming, it may not be solely responsible for the full acquisition of metastatic phenotypes. Our results using lipofection-mediated delivery of recombinant Cdt showed that, although DNA damage was induced at comparable concentrations, an increase in migration required substantially higher doses of Cdt than those estimated in Aa-EVs did. These findings imply that additional components within Aa-EVs act as enhancers that cooperate with Cdt to efficiently induce EMT and metastatic features. Identifying such cofactors within Aa-EVs will be important for future investigations.

Interestingly, the human-specific nature of this mechanism and its dependence on EGFR signaling suggest that the oncogenic potential of Aa-EVs may vary among cancer types because of their reliance on EGFR signaling. Indeed, a recent study reported that Aa-EVs exerted antitumorigenic rather than prometastatic effects in oral squamous cell carcinoma (*50*). This apparent discrepancy may reflect tissue-specific differences in cellular context, EGFR dependency, and the balance between DDR-induced cytostasis and SASP-driven tumor promotion. Such context-dependent outcomes underscore the complexity of host–microbe interactions in cancer biology.

In the present study, we focused primarily on the effects of Aa-EVs on established tumor cells. However, EVs derived from oral pathogens are likely to influence multiple components of the tumor microenvironment. For instance, EVs may modulate tumor-associated macrophages, cancer-associated fibroblasts, and endothelial cells, thereby altering the immune and stromal landscapes that support tumor growth and metastasis (*3*, *7*, *51*, *52*). Given that SASP-like cytokines such as IL-1β, IL-6, and CXCL8 were upregulated in Aa-EV-treated PAC cells, these cells might, in turn, remodel the TME through paracrine signaling. Future investigations should therefore explore how Aa-EVs affect the crosstalk between PAC cells and nonmalignant cells within the TME.

Interestingly, only Aa-EVs, but not Pg- or Fn-derived EVs, enhanced the migration and invasion of PAC cells in our study. However, previous epidemiological and clinical studies have implicated Pg and Fn in the development of PAC (*12*, *13*). It is conceivable that their EVs participate in distinct phases of the tumorigenic cascade; for example, Pg-and Fn-derived EVs might contribute to tumor initiation or promotion, whereas Aa-EVs drive tumor progression and metastasis. This sequential involvement is consistent with the classical multistep model of cancer development (initiation, promotion, and progression) (*53*). Comprehensive comparative analyses of periodontopathic bacterial EVs are needed to delineate their cooperative or sequential roles in the oral–pancreatic cancer axis.

In summary, this study identifies a previously unrecognized mechanism by which Aa-derived EVs promote the metastatic features of pancreatic cancer cells through Cdt-mediated DNA damage and subsequent activation of the EGFR–ERK/Akt signaling axis. Our data also suggest that Aa-EV-induced senescence-like states may represent a transitional step toward malignant transformation, potentiated by EV-associated enhancers that amplify Cdt activity. These findings establish a new conceptual framework linking oral bacterial EVs to pancreatic cancer progression and provide mechanistic insight into the epidemiological connection between periodontitis and pancreatic cancer.

Our findings have several limitations. Because of the species-specific activity of Cdt, in vivo validation using conventional rodent models was not feasible. Consequently, the current study was restricted to cancer cell–autonomous effects and did not address complex interactions with immune or stromal cells in the TME. Moreover, although Aa-EVs contain multiple bacterial components, we focused primarily on Cdt; however, other EV cargoes may also contribute to the observed phenotypes. Future studies should therefore employ advanced in vitro coculture systems and humanized in vivo models to dissect the contributions of these components and to determine whether chronic exposure to Aa-EVs can accelerate pancreatic cancer progression or contribute to its initiation.

## Materials and Methods

### Bacterial strains and culture conditions

Three periodontopathic bacterial strains—*Porphyromonas gingivalis*, *Fusobacterium nucleatum*, and *Aggregatibacter actinomycetemcomitans*—were used in this study. *P. gingivalis* strain ATCC 33277 was grown in brain heart infusion (BHI) broth (BD) supplemented with 5 µg/mL hemin and 1 µg/mL menadione (BHI-HM) or on BHI-HM blood agar plates. *F. nucleatum* strain #20 and all *A. actinomycetemcomitans* strains, including the D7SS strain used for genetic manipulation, were grown in BHI broth or on BHI blood agar. All peridontopathic bacteria were grown in an anaerobic chamber (MiniMACS anaerobic workstation, Don Whitley Scientific Ltd.) in 80% N_2_, 10% H_2_, and 10% CO_2_ at 37°C, as described previously (*54*, *55*). For gene manipulation of A. actinomycetemcomitans, cells were incubated and selected on BHI supplemented with 5% heat-inactivated horse serum (Japan Bioserum) (BHI-HS agar) supplemented with appropriate antibiotics in a humidified atmosphere of 5% CO_2_ at 37°C. *Escherichia coli* strains DH5α and BL21(DE3) were also used for cloning and recombinant protein expression, respectively. E. coli was grown in Luria–Bertani (LB) broth (BD) or on LB agar plates under aerobic conditions at 37°C. To select transformants, ampicillin (100 µg/mL) or kanamycin (25 µg/mL) was added to the medium as appropriate.

### Cell culture

The human pancreatic cancer cell lines PANC-1 (RIKEN BRC #RCB2095) and SUIT-2 (JCRB Cell Bank #JCRB1094) and the murine pancreatic cancer cell line KPC, kindly provided by Dr. Yoshihiro Nishikawa, were used (*27*). PANC-1 and SUIT-2 cells were cultured in RPMI-1640 medium, and KPC cells were cultured in high-glucose DMEM supplemented with 10% fetal bovine serum, 100 U/mL penicillin, and 100 µg/mL streptomycin in a humidified atmosphere of 5% CO_2_ at 37°C. Cells were treated

### Isolation of bacterial extracellular vesicles

Bacterial EVs were isolated by ultracentrifugation as previously described (*54–56*). Briefly, bacterial culture supernatants were collected by centrifugation at 15,000 × g for 20 min at 4°C and filtered through 0.45 µm and 0.22 µm PVDF membranes (Stericup, Merck #S2HVU01RE and #S2GVU01RE, respectively). The filtrates were ultracentrifuged at 100,000 × g for 1.5 h at 4°C to pellet the EVs, which were subsequently resuspended in PBS and stored at -30°C until use. Protein concentrations were determined by a Bradford assay (Bio-Rad #5000006). EV morphology was analyzed by field-emission scanning electron microscopy (FE-SEM) (*56*), and the size distribution and particle number were quantified by a NanoFCM system (*56*).

### Uptake of EVs by host cells

EVs were fluorescently labeled with CellTrace™ Far Red (Invitrogen #34564). Briefly, Aa-EVs (10 µg/mL) were incubated with 10 µM dye for 30 min at room temperature and ultracentrifuged twice at 100,000 × g for 1.5 h at 4°C to remove free dye. The labeled EVs were resuspended in PBS and stored at 4°C. Fluorescently labeled Aa-EVs were incubated with PANC-1 or KPC cells for 2 h. After fixation with paraformaldehyde and staining with phalloidin (Proteintech #PF00001) and DAPI, images were captured using an EVOS M7000 microscope (Thermo Fisher Scientific). Fluorescence images were linearly adjusted for brightness and contrast equally across the entire image using Adobe Photoshop, with identical parameters applied to all samples, including controls.

### DNA damage assay

DNA damage was evaluated using a DNA damage detection kit (Dojindo #G265) according to the manufacturer’s instructions. After 24 h of Aa-EV treatment, the cells were fixed and stained with γH2AX, phalloidin, and DAPI. γH2AX foci were imaged using a BZ-X810 microscope (Keyence). Fluorescence images were linearly adjusted for brightness and contrast equally across the entire image using Adobe Photoshop, with identical parameters applied to all samples, including controls.

### Construction of the cdtABC knockout and complement strains of Aa

Gene manipulation was performed using the *A. actinomycetemcomitans* strain D7SS as previously described (*40*). To generate the *cdtABC* knockout strain (Cdt-KO), approximately 1-kb fragments flanking the *cdtABC* operon were amplified from D7SS genomic DNA using the primer pairs listed in Table S1, and a kanamycin resistance cassette was amplified from the plasmid pMV261 (*57*). These fragments were assembled into NdeI-digested pGEM-T Easy (Promega #A1360) using NEBuilder^®^ HiFi DNA Assembly Master Mix (New England Biolabs #E2621), yielding the plasmid pTYAa01 (Fig. S11A). The linearized plasmid (SpeI-digested) was introduced into the D7SS by natural transformation, and transformants were selected on BHI-HS agar supplemented with 25 µg/mL kanamycin to obtain the Δ*cdtABC*::KmR mutant.

Next, the kanamycin resistance cassette in the mutant genome was replaced by a spectinomycin resistance cassette (SpcR) to generate Δ*cdtABC*::SpcR. Approximately 1-kb fragments flanking the km cassette and the spectinomycin-resistance cassette (amplified from pUTmini-Tn5 Sm) were assembled into NdeI-digested pGEM-T Easy (pTYAa02; Fig. S11B) using NEBuilder. The linearized pTYAa02 plasmid (SpeI-digested) was introduced into the Δ*cdtABC*::KmR strain by natural transformation, and transformants were selected on BHI-HS agar supplemented with 50 µg/mL spectinomycin. Correct recombination and cassette replacement were confirmed by PCR. To construct the complemented strain (Δ*cdtABC*::*cdtABC*-His₆::Spcᴿ), each subunit of the *cdtABC* operon (*cdtA*, *cdtB*, and *cdtC*) was first amplified from D7SS genomic DNA with a C-terminal His₆ tag, generating *cdtA*-His₆, *cdtB*-His₆, and *cdtC*-His₆ fragments. Each fragment was individually cloned and inserted into pGEM-T Easy, yielding the pTYAa03–05 plasmid (Figs. S11, C to E). To construct the complementation vector, pTYAa02 was linearized by PCR at the junction between the spectinomycin resistance cassette and the downstream flanking region, and three fragments (*cdtA*-His₆, *cdtB*-His₆, and *cdtC*-His₆), including the native promoter and 5′ untranslated region (UTR), were assembled into the linearized pTYAa02 backbone using NEBuilder HiFi DNA Assembly, yielding pTYAa06 (Fig. S11F). Linearized pTYAa06 was introduced into the Δ*cdtABC*::Kmᴿ strain by natural transformation, and transformants were selected on BHI-HS agar supplemented with 50 µg/mL spectinomycin. Correct replacement of the km cassette with the *cdtABC*-His₆::Spcᴿ locus was verified by PCR, and the expression of His₆-tagged Cdt proteins was confirmed by Western blotting using an anti-His antibody (MBL).

### Purification of recombinant CdtA, CdtB, and CdtC

Coding sequences of *cdtA*, *cdtB*, and *cdtC* lacking their native signal peptides were amplified from plasmids pTYAa03–05, which encode C-terminal His₆-tagged versions of each subunit. The amplified fragments were digested and cloned, and inserted into pET22b(+) (Novagen) as follows: *cdtA* and *cdtB* using NdeI/XhoI (resulting in pTYAa07 and pTYAa08), and *cdtC* using NcoI/XhoI (resulting in pTYAa09) (see Fig. S12). The His₆-tag sequence encoded by pET22b(+) was not used.

For soluble expression, CdtA and CdtB were coexpressed with molecular chaperones using the plasmids pKJE7 and pG-KJE8 (Takara Bio), respectively. Protein expression was induced by 1 mM isopropyl β-D-1-thiogalactopyranoside (IPTG) at 37°C for 4 h. Cells expressing CdtA or CdtB were harvested and lysed by sonication in buffer (50 mM Tris-HCl (pH 8.0), 300 mM NaCl, and 10 mM imidazole). Recombinant proteins were purified using HisTrap™ HP columns (Cytiva) following the manufacturer’s instructions. To remove residual chaperones from the CdtA fractions, an additional casein–ATP wash was performed.

CdtC was expressed under the same induction conditions and purified from the periplasmic fraction obtained by osmotic shock, followed by HisTrap™ HP affinity chromatography. Protein purity and molecular weights were confirmed by SDS–PAGE and Coomassie Brilliant Blue staining (see Fig. S5).

For treatment of PANC-1 cells with recombinant Cdt (rCdt), purified CdtA, CdtB, and CdtC were mixed at equimolar ratios. Protein transfection was performed using Pro-DeliverIN (OZ Biosciences #PI10100) according to the manufacturer’s instructions to assess the effects of intracellular delivery of rCdt.

### Wound healing assay

PANC-1 and KPC cells were seeded on 24-well plates (5×10^4^ – 1×10^5^ cells per well) and incubated until they reached 80–90% confluence. After the medium was replaced with fresh medium, the EVs from periodontopathic bacteria were treated at concentrations ranging from 10–1000 ng/mL. After 24 h of EV treatment, the cells were scratched using 200 µL micropipette tips. The cells were then washed with PBS twice and treated with the EVs again after the culture medium was replaced. For inhibition of the EGFR signaling pathway, an ERK1/2 inhibitor SCH772984 (Cayman #19166) and/or an Akt inhibitor MK-2206 (Selleck #S1078), or a pan-EGFR inhibitor sapitinib (AZD8931) (Selleck #2192) were used. Microscope images were captured using a BZ-X800 (Keyence), and the open area was quantified using ImageJ.

### Transwell invasion assay

PANC-1 or SUIT-2 cells were seeded on 6-well plates (5×10^5^ cells per well) and incubated for 24 h. Then, the cells were treated with EVs after the medium was replaced with fresh complete RPMI. After 48 h of EV treatment, the cells were trypsinized and harvested. A total of 2×10^4^ cells, which were suspended in RPMI-1640 supplemented with 200 µL of 0.1% FBS, were seeded on the upper side of Transwell^®^-clear inserts with a diameter of 6.5 mm and 8 µm pores (Corning), which had been coated with type I collagen (Koken) on the bottom side. In addition, 500 µL of complete RPMI was added to the bottom space of the inserts, and the cells were incubated for 14 h. The cells were fixed with neutral formalin, permeabilized with methanol, and then stained with 0.1% crystal violet. The number of invaded cells on the bottom side of the inserts was determined by ImageJ. The experiments were performed in triplicate and repeated twice independently.

### Western blotting

To evaluate EMT, PANC-1 cells were treated with EVs (1,000 ng/mL) for 48 h, followed by incubation in EV-free complete RPMI for an additional 12 h. To assess hyperactivation of the EGFR or TGFβ signaling pathway, cells were treated with EVs (1,000 ng/mL) for 48 h, followed by incubation in serum-free RPMI for 3 h. Cells were then stimulated with human EGF (10 ng/mL; Fujifilm Wako Pure Chemical #058-09521) or human TGFβ1 (10 ng/mL; Fujifilm Wako Pure Chemical #209-21571). Cells were lysed in lysis buffer (20 mM Tris-HCl (pH 7.5), 2 mM EDTA, 150 mM NaCl, 1% (v/v) Nonidet™ P-40, 1 mM PMSF, and 1x protease inhibitor cocktail (Nacalai Tesque #25955-11)). The proteins were resolved by sodium dodecyl sulfate–polyacrylamide gel electrophoresis and transferred to Immobilon-P PVDF membranes (Millipore #IPVH00010). Membranes were probed with primary antibodies against N-cadherin (Cell Signaling Technology #13116), E-cadherin (Cell Signaling Technology #3195), Vimentin (Cell Signaling Technology, #5741), phospho-Smad2/3 (Cell Signaling Technology #8828), phospho-EGFR (Proteintech #30278-1-AP), phospho-ERK1/2 (Proteintech #28733-1-AP), total ERK1/2 (Proteintech #11257-1-AP), phospho-Akt (Proteintech #66444-1-Ig), total Akt (Proteintech #7074), or His-tag (MBL #D291-7). Membranes were then incubated with horseradish peroxidase-conjugated anti-mouse IgG (Proteintech RGAM601) or anti-rabbit IgG (Cell Signaling Technology #7074) secondary antibodies (1:5,000). Protein signals were visualized using ImmunoStar^®^ Zeta or ImmunoStar^®^ LD regents (Fujifilm Wako Pure Chemical) and detected with a FUSION Solo S imaging system (Vilber). Band intensities were quantified using ImageJ software (NIH).

### RNA isolation

PANC-1 cells were treated with EVs (1,000 ng/mL) for 48 h and subsequently incubated with EV-free complete RPMI for 3 h. Total RNA from OMV-treated cells was isolated using TRI reagent^®^ (Molecular Research Center #TR118) and a DirectZol™ Microprep kit (Zymo Research #R2062) as described previously (*56*). Afterward, the quality of the RNA was evaluated using a NanoDrop 2000 (Thermo Scientific). For qPCR, cDNA was synthesized using ReverTraAce^®^ qPCR RT Master Mix with gDNA Remover (Toyobo #FSQ-301). Gene expression levels were quantified with Luna^®^ Universal qPCR Master Mix (New England BioLabs #M3003). The sequences of the primer sets used in this study are listed in Table S1. PCR was performed on a LightCycler 96 instrument (Roche). For data normalization, the relative expression levels of target genes were calculated via the comparative CT (ΔΔCT) method and compared with those of 18S rRNA as internal standards. The experiments were performed independently three times.

### RNA sequencing and data processing

Total RNA, isolated as described above, was submitted to KOTAI Biotechnologies Inc. (Osaka, Japan) for sequencing. RNA integrity was assessed using a Bioanalyzer 2100 (Agilent Technologies). Library preparation was performed with the NEBNext^®^ Ultra™ II Directional RNA Library Prep Kit (New England Biolabs) according to the manufacturer’s instructions, and sequencing was conducted on the DNBSEQ-T7 platform (MGI Tech) using a 150 bp paired-end strategy.

The raw sequencing data were quality checked using FastQC and aligned to the H. sapiens reference genome (GRCh38) using STAR. Gene-level quantification was performed using RSEM to generate raw count data. Primary processing was conducted using the nf-core/rnaseq workflow (v3.14.0) implemented in Nextflow v23.10.1 (*58–60*), ensuring full reproducibility through Singularity containers.

Subsequent analyses were performed using RNAseqChef (https://imeg-ku.shinyapps.io/RNAseqChef/) (*61*). Normalization and differential expression analyses were conducted using DESeq2. Hallmark pathway enrichment analysis of upregulated genes was performed based on the hallmark gene sets from the Molecular Signatures Database (MSigDB v2024.1.Hs) (*62*). Gene set enrichment analysis (GSEA) was performed using GSEA software (v4.4.0, Broad Institute) (*63*) with the MSigDB Hallmark gene sets (*62*). We used ChatGPT to assist with R and Python script development.

### Clinical data acquisition and analysis

Clinical prognosis and transcriptomic data of patients with pancreatic cancer (PAC; n = 135) and colorectal cancer (CRC; n = 110) were obtained from the Genomic Data Commons (GDC) Data Portal (https://portal.gdc.cancer.gov). Transcriptome profiles were compared between patients with a complete response (CR) and those with progressive disease (PD). Raw count data were normalized using the DESeq2 package (v1.44.0) (*64*) in R (v4.4.1). GSEA was performed using the clusterProfiler package (v4.12.6) (*65*) with MSigDB hallmark gene sets (v24.1.0) (*62*), and gene set variation analysis (GSVA) was conducted using the GSVA package (v1.52.3) (*66*) to estimate the enrichment scores of EMT-, p53-, and E2F-related pathways in PAC patients. Survival curves were generated using the Kaplan–Meier method, and differences were assessed by the log-rank test. We used ChatGPT to assist with R and Python script development.

### Cell viability and proliferation assays

For the colorimetric cell viability assay, PANC-1 cells or SUIT-2 cells were plated at 5 × 10^3^ or 1 × 10^4^ cells/well, respectively, in a 96-well plate. The next day, after the medium was replaced with fresh culture medium, the cells were treated with bacterial EVs at a concentration of 1 µg/mL. After 48 h of EV treatment, cell viability was quantified using a Cell Counting Kit-8 (Dojindo #CK04) according to the manufacturer’s instructions. For the time-lapse cell proliferation assay, PANC-1 cells or SUIT-2 cells were plated at 2.5 × 10^4^ or 5 × 10^4^ cells/well, respectively, in a 24-well plate. The next day, the cells were treated with EVs derived from the WT and *cdtABC* mutant Aa strains. Microscopy images were captured using the CM30 Cell Culture Monitoring System (EVIDENT) every 3 h in 4 areas per well. The number of cells in each well was calculated as the average of the numbers of cells in each of the areas.

### Statistical analysis

All the quantitative data are presented as the mean ± SEM. Statistical analyses were performed using GraphPad Prism. One-way ANOVA followed by Tukey’s HSD or unpaired t tests were used as appropriate. Differences were considered statistically significant at P < 0.05.

## Acknowledgments

We thank Tadaki Suzuki, Akira Ainai, Hirotaka Kobayashi, and Fumiko Takashima (National Institute of Infectious Diseases, Japan), as well as Yusuke Kouda and Naoki Narisawa (Nihon University, Japan), for their technical support. We are also grateful to members of the Research Support Platform of Osaka Metropolitan University Graduate School of Medicine. We sincerely thank Jan Oscarsson (Umeå University, Sweden) for providing the *A. actionomycetemcomitans* D7SS strain; Mariko Naito and Hideaki Hayashida (Nagasaki University, Japan) for providing the *A. actinomycetemcomitans ltxA* mutant and its parent strain; and Yoshihiro Nishikawa (Kyoto University, Japan) for providing the murine KPC cells. Illustrations were created using BioRender. During the preparation of this manuscript, the authors used ChatGPT and Grammarly to assist with English editing.

## Funding

This work was conducted with the support of the following funding.

The Japanese Ministry of Education, Culture, Sports, Science and Technology (MEXT) (24K13232, to TY)

The Japanese MEXT (21K18284, to RN) The Japanese MEXT (21KK0164, to RN)

The Takeda Science Foundation (2024042293, to TY) The SENSHIN Medical Research Foundation (to TY).

## Author contributions

Conceptualization: TY, RN

Methodology: TY, MS

Investigation: TY, MS, KA

Visualization: TY, RN, MS

Supervision: RN, MS, HA

Writing—original draft: TY, MS

Writing—review & editing: TY, RN, MS, KA, HA

## Data and materials availability

All data are available in the main text or the supplementary materials. RNA-seq data have been deposited in the DDBJ Sequence Read Archive (DRA), and the Genomic Expression Archive (GEA) under the accession numbers DRA025841 and E-GEAD-1185, respectively. These data will be made publicly available upon publication. All the code and analysis pipelines are available upon reasonable request. The plasmids and bacterial strains generated in this study are available upon request to the Lead Contact (Takehiro Yamaguchi).

---

**Fig. S1.**
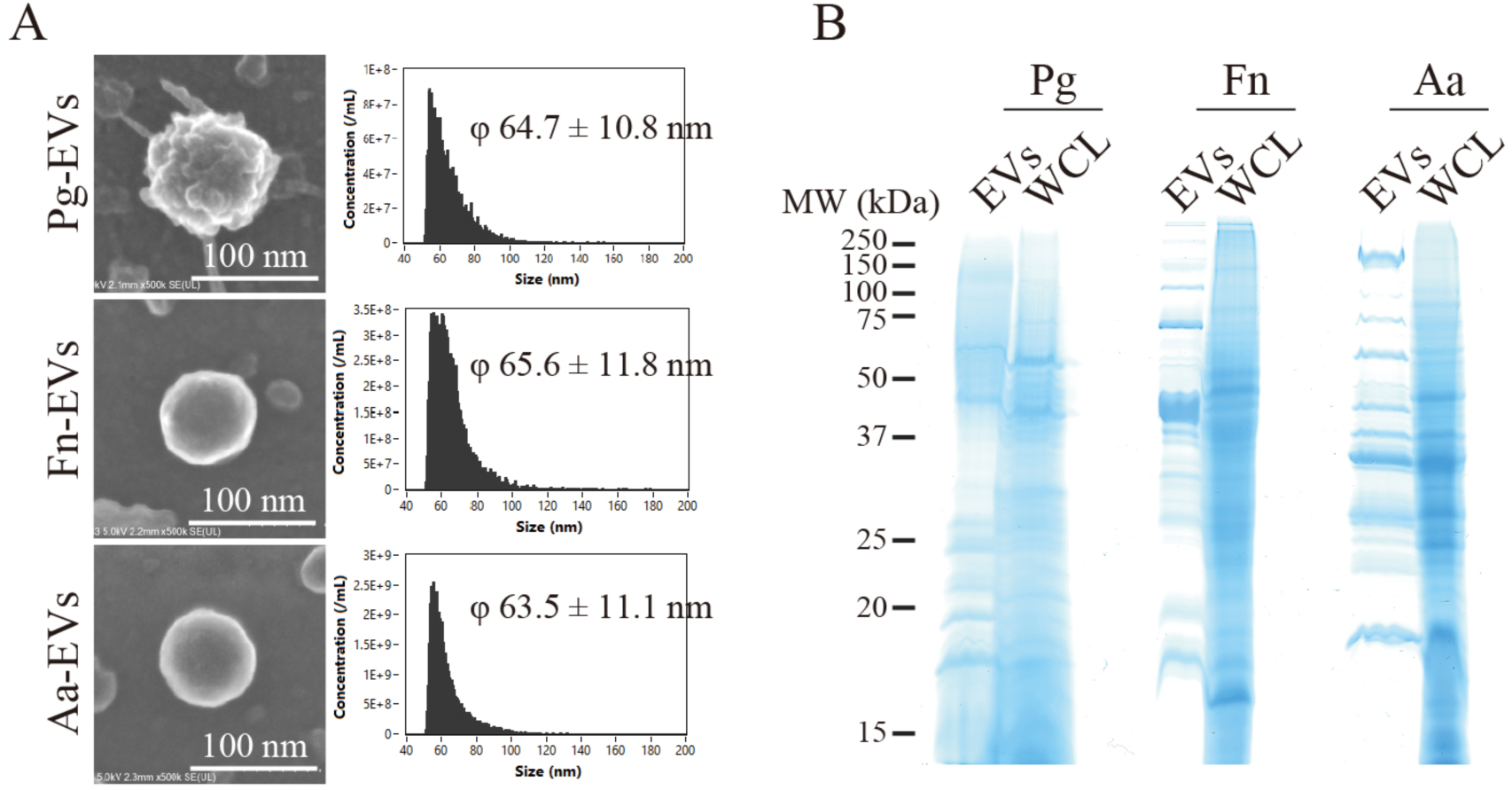
Analysis of the morphology and protein expression of EVs derived from pancreatic cancer-associated periodontopathic bacteria. **(A)** Morphological analysis of EVs derived from *Porphyromonas gingivalis* (Pg-EVs), *Fusobacterium nucleatum* (Fn-EVs), and *Aggregatibacter actinomycetemcomitans* (Aa-EVs) analyzed by field emission–scanning electron microscopy (left) and nanoflow cytometry (right). **(B)** Protein expression profiles of EVs and whole-cell lysates (WCLs) of donor bacteria were analyzed by SDS–PAGE.

**Fig. S2.**
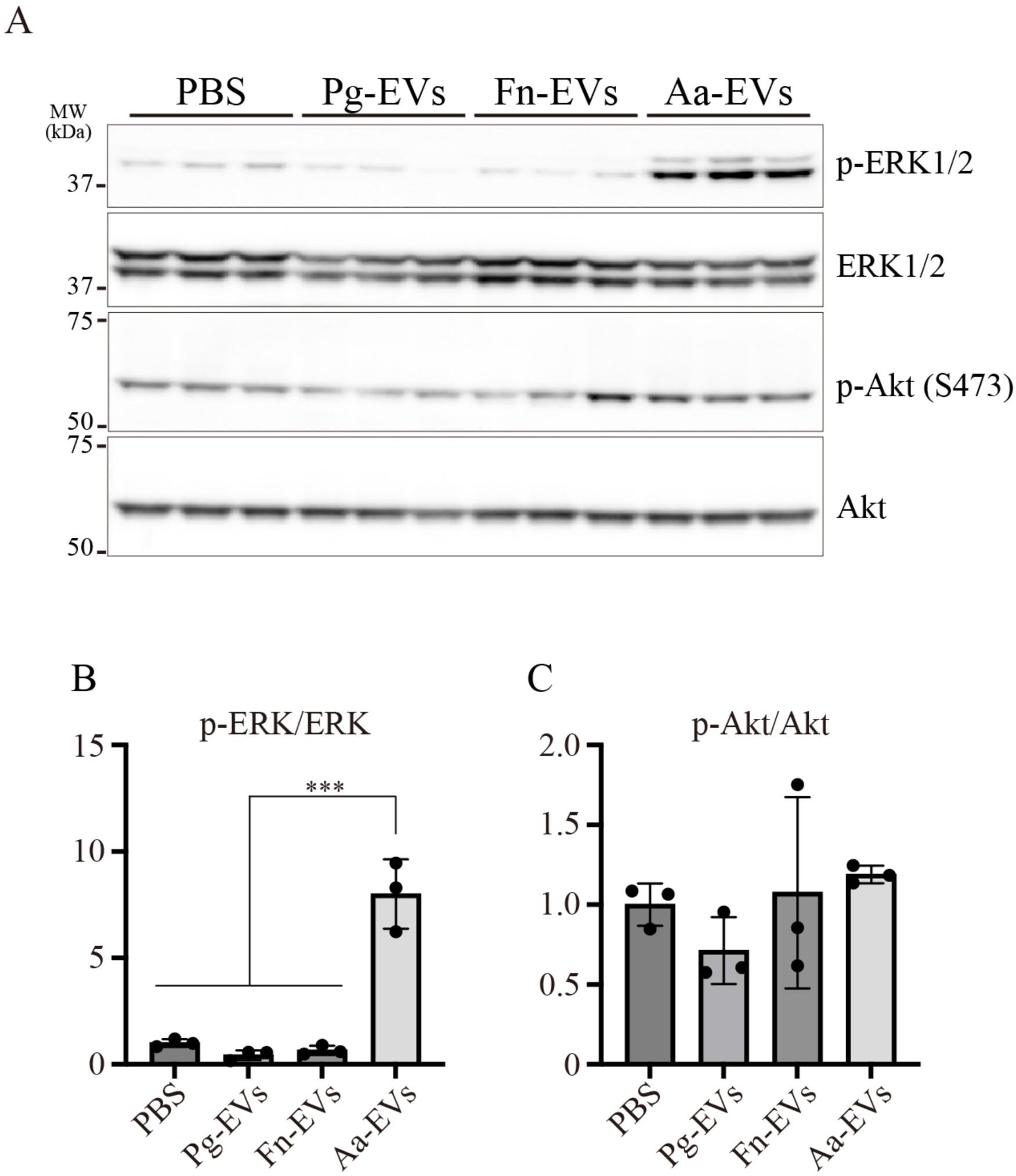
ERK1/2 and Akt activation in PANC-1 cells treated with periodontopathic bacterial EVs. **(A)** Western blot analysis of ERK1/2 and Akt in PANC-1 cells treated with EVs derived from periodontopathic bacteria. **(B)** Quantification of phosphorylated ERK1/2 (p-ERK) relative to total ERK1/2. **(C)** Quantification of phosphorylated Akt (S473) relative to total Akt. The data are presented as the means ± SDs. *P < 0.05; **P < 0.01; ***P < 0.001 (Tukey’s test).

**Fig. S3.**
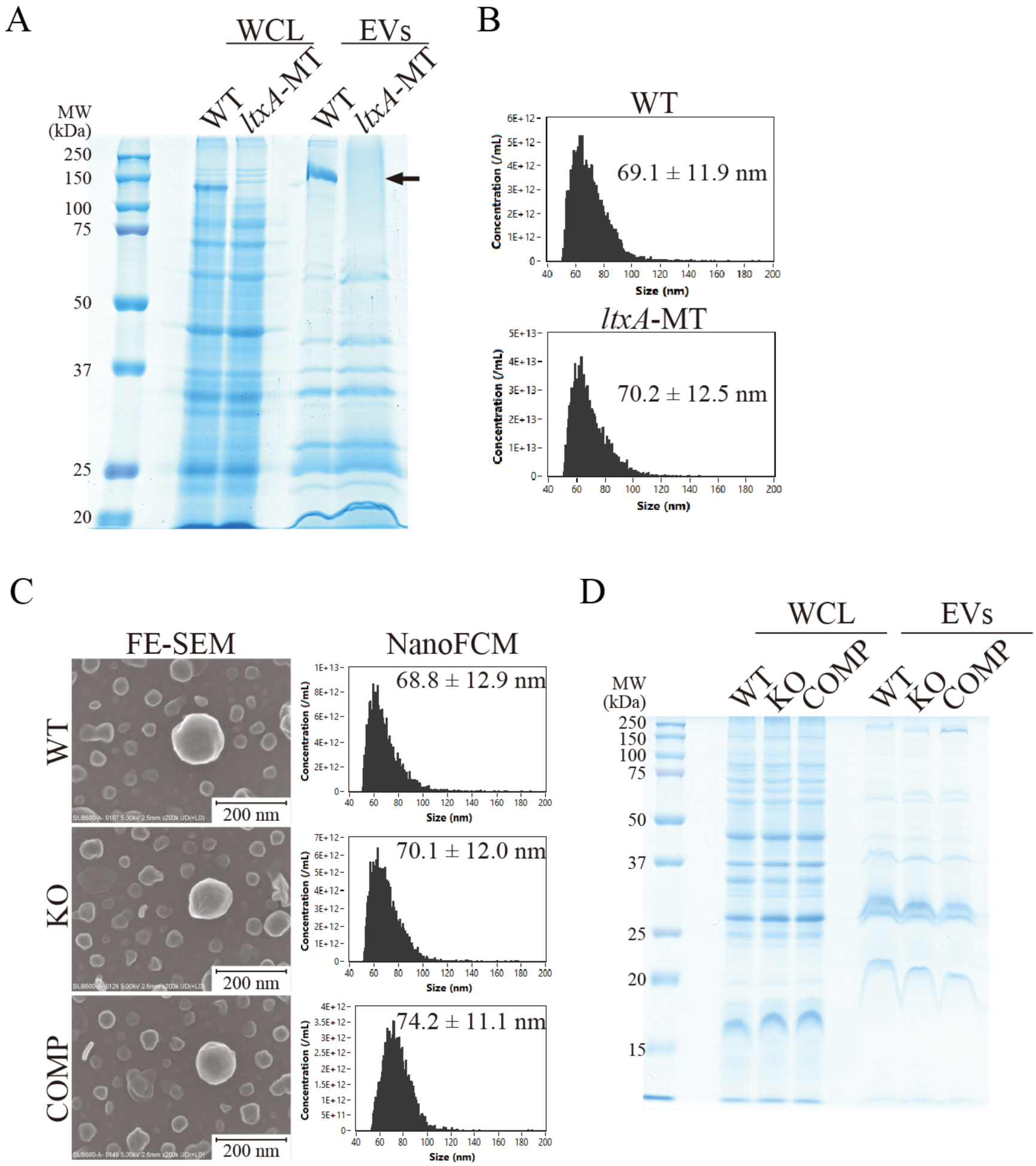
Characterization of EVs derived from genetically engineered *A. actinomycetemcomitans* strains. (A, **B)** Protein expression (A) and size distribution (B) of EVs derived from wild-type (WT) *A. actinomycetemcomitans* JP2 and its *ltxA* mutant (*ltxA*-MT). **(C)** Morphological analysis (left) and size distribution (right) of EVs derived from WT, *cdtABC* knockout (KO), and *cdtABC* complement (COMP) *A. actinomycetemcomitans* D7SS strains. **(D)** Protein expression profiles of EVs derived from WT, KO, and COMP *A. actinomycetemcomitans* D7SS strains.

**Fig. S4.**
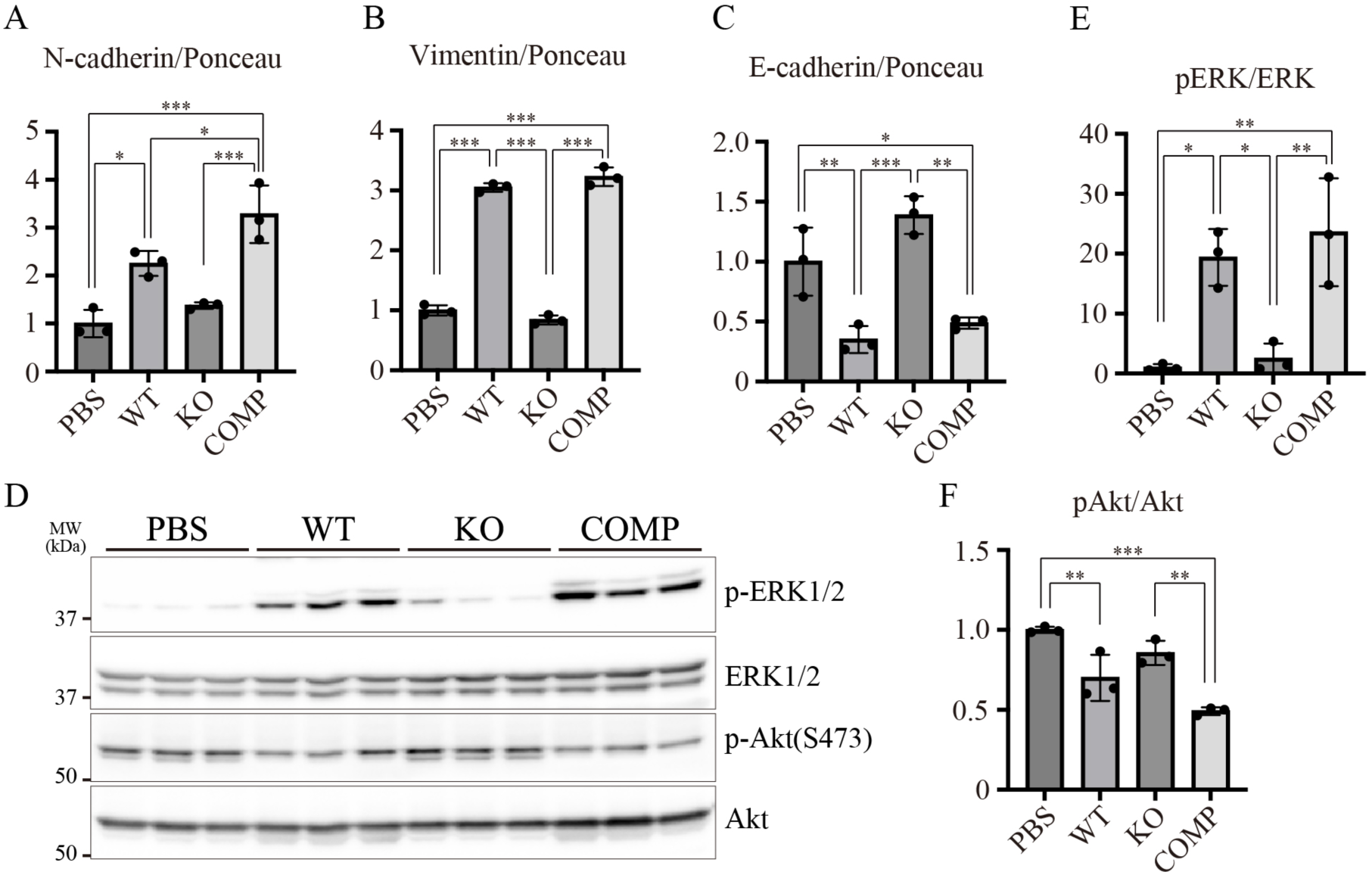
Cdt-dependent induction of epithelial–mesenchymal transition in PANC-1 cells. (**A**–**C**) Quantification of the mesenchymal markers N-cadherin (A) and vimentin (B) and the epithelial marker E-cadherin (C), corresponding to Fig. 2F. (**D**–**F**) Evaluation of Cdt-dependent activation of ERK1/2 and Akt signaling (D). Quantification of phosphorylated ERK1/2 relative to total ERK1/2 (E) and phosphorylated Akt (S473) relative to total Akt (F). The data are presented as the means ± SDs. *P < 0.05; **P < 0.01; ***P < 0.001 (Tukey’s test).

**Fig.S5.**
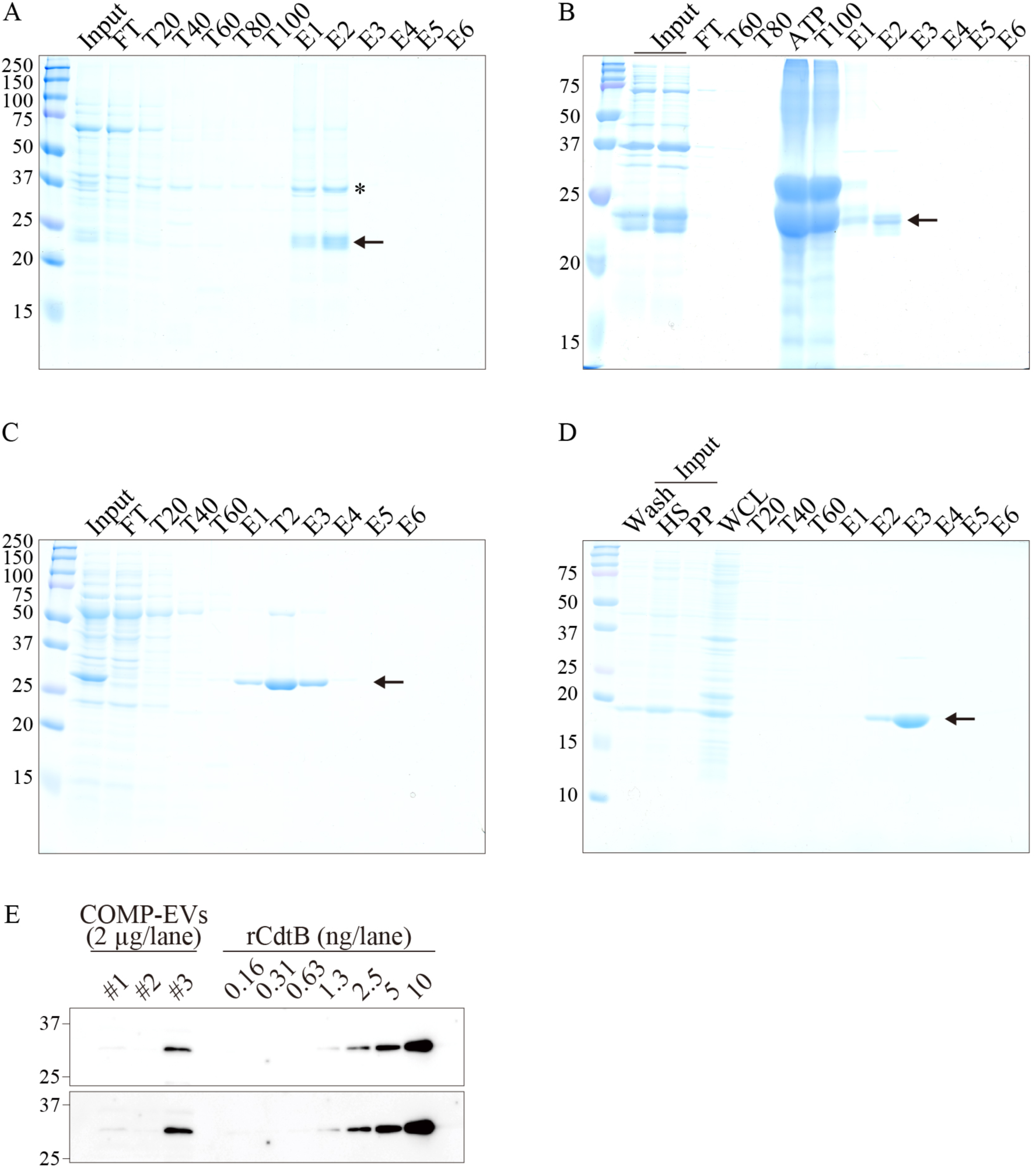
Purification of recombinant CdtA, CdtB, and CdtC and quantification of the CdtB content in His_6_-tagged Cdt-expressing Aa-EVs. **(A, B)** Two-step purification of recombinant CdtA from *E. coli* BL21 (DE3) coexpressing His_6_-tagged CdtA and the chaperones DnaK, DnaJ, and GrpE. (A) First Ni–NTA purification. (B) Second purification using casein–ATP washes to remove chaperones. **(C)** Purification of recombinant CdtB coexpressed with GroES, GroEL, DnaK, DnaJ, and GrpE. **(D)** Purification of recombinant CdtC from *E. coli* BL21 (DE3) expressing His_6_-tagged CdtC. **(E)** Quantification of His_6_-tagged CdtB content in *cdtABC* complement Aa-derived EVs by immunoblotting.

**Fig. S6.**
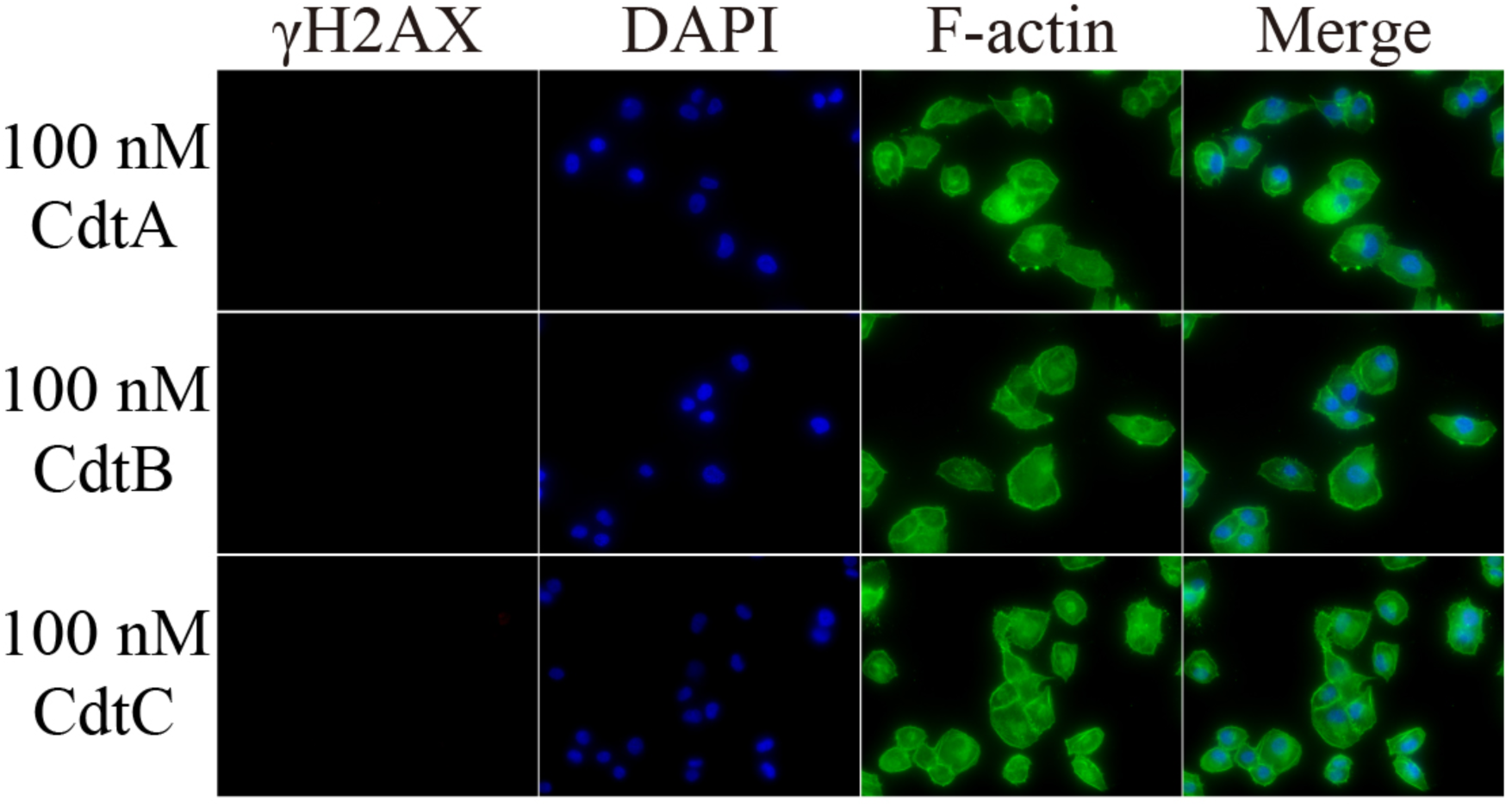
γΗ2ΑΧ staining of PANC-1 cells treated with recombinant Cdt subunits. Representative immunofluorescence images of γH2AX (red), F-actin (green), and nuclei (blue) in PANC-1 cells treated with recombinant CdtA, CdtB, or CdtC individually.

**Fig. S7.**
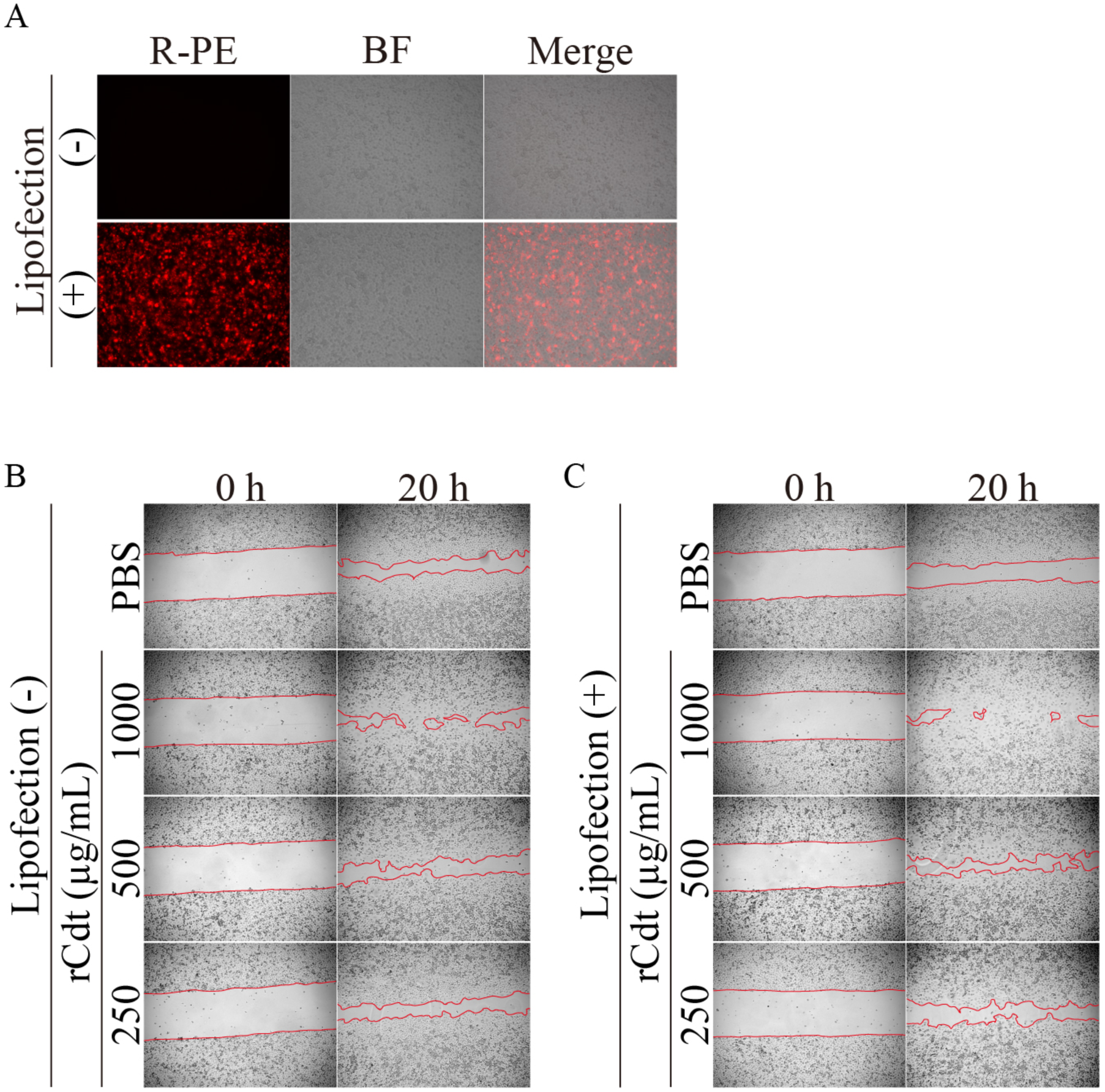
Effect of rCdt transfection on cellular migration (related to Figs. 4, J to L). **(A)** Confirmation of successful protein transfection using R-phycoerythrin (R-PE). **(B, C)** Microscopy images of the wound healing assay in rCdt-treated PANC-1 cells with or without lipofection reagent.

**Fig. S8.**
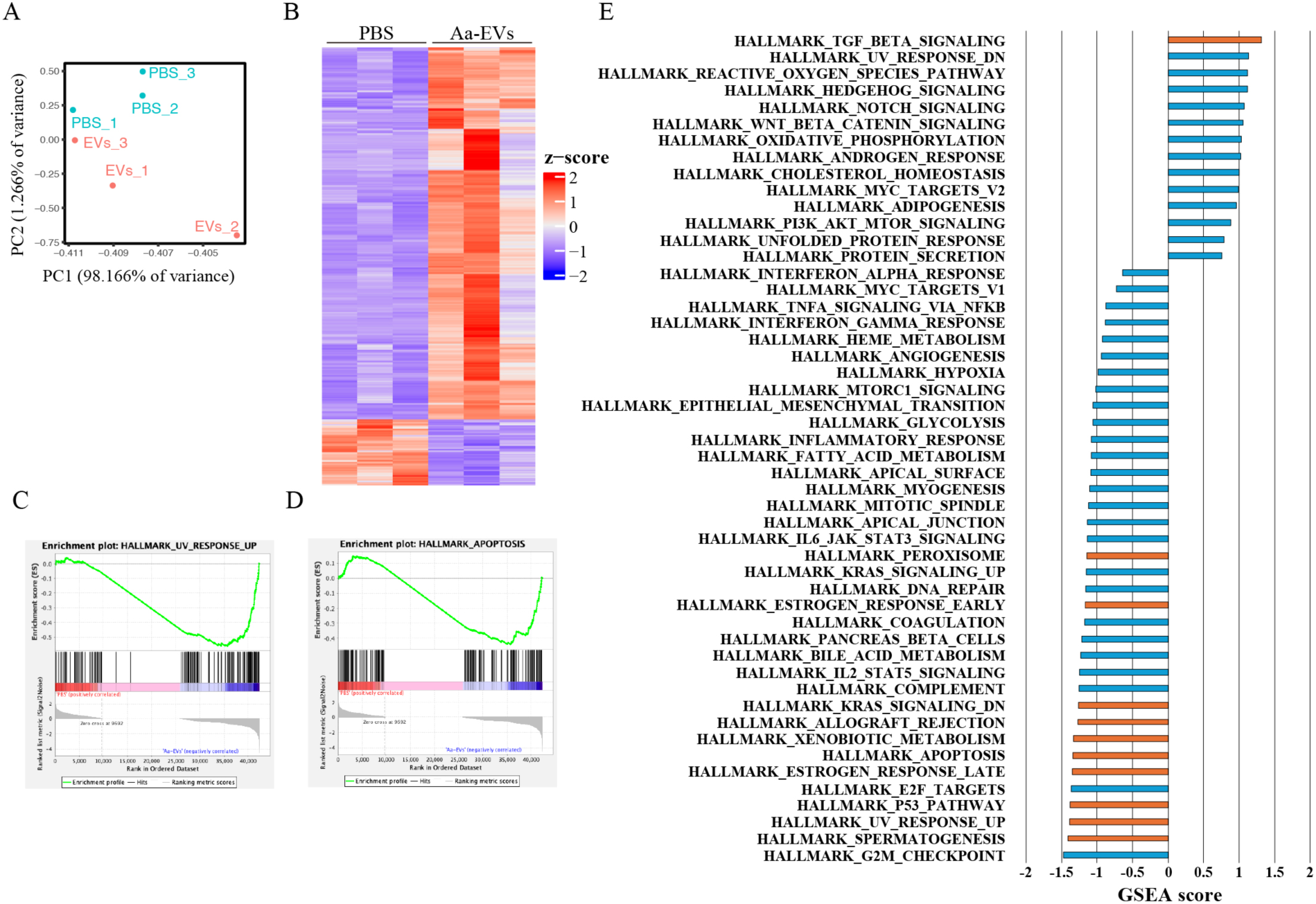
Transcriptomic analysis of Aa-EV-treated PANC-1 cells. **(A)** Principal component analysis using transcriptomic data from PBS-treated and Aa-EV-treated PANC-1 cells. **(B)** Heatmap showing representative results of differential expression analysis. **(C-E)** Gene set enrichment analysis (GSEA) comparing PBS-treated PANC-1 cells and Aa-EV-treated PANC-1 cells. (C) GSEA plot for the UV response gene set. (D) GSEA plot for the apoptosis gene set. (E) Bar plot summarizing the results of GSEA using the hallmark gene sets from the Molecular Signature Database (MSigDB); pathways showing significant enrichment are highlighted in orange.

**Fig. S9.**
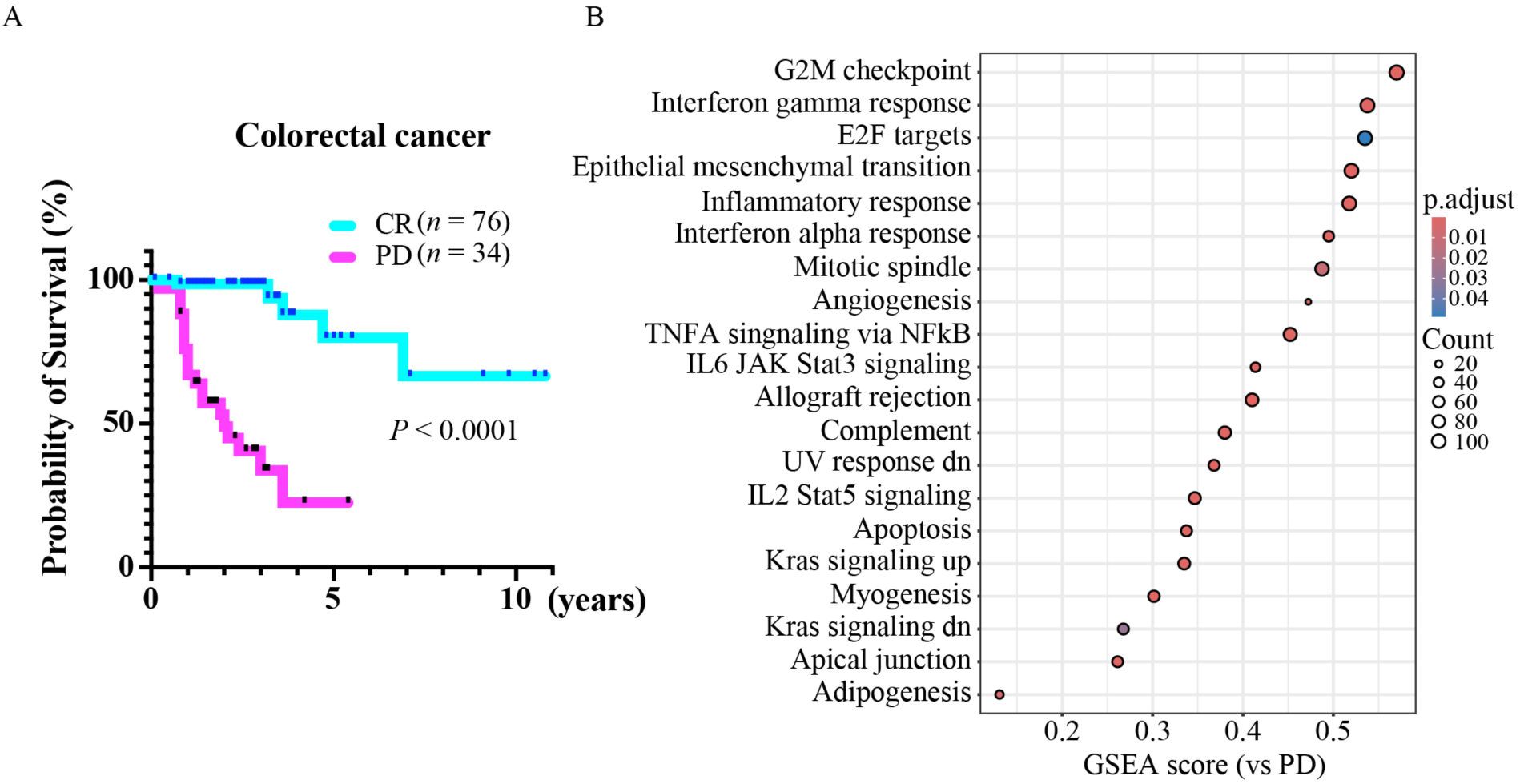
Transcriptomic analysis of cancer tissues was performed using data from the GDC Data Portal. **(A)** Kaplan–Meier survival analysis of colorectal cancer (CRC) patients with complete response (CR) or progressive disease (PD). **(B)** GSEA results comparing CR versus PD in CRC patients.

**Fig. S10.**
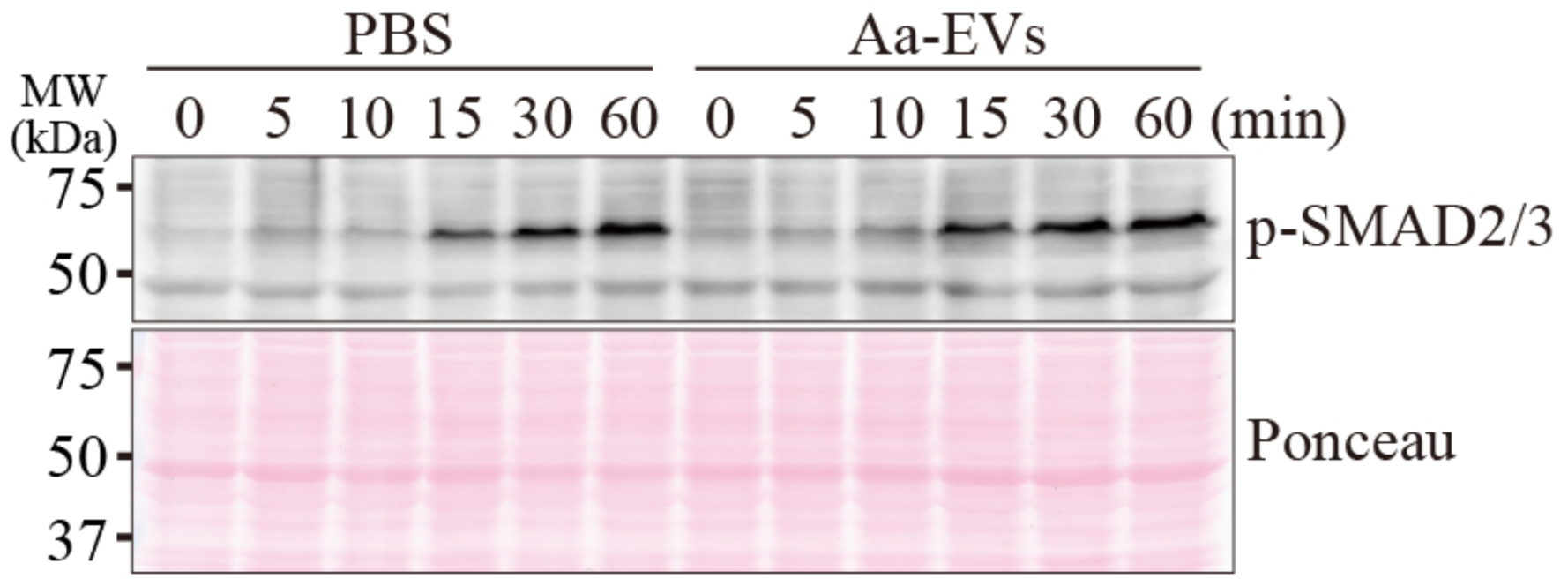
Aa-EV pretreatment did not enhance TGFβ signaling in PANC-1 cells. Comparison of TGFβ signaling activation in TGFβ1-treated PANC-1 cells with or without Aa-EV pretreatment.

**Fig. S11.**
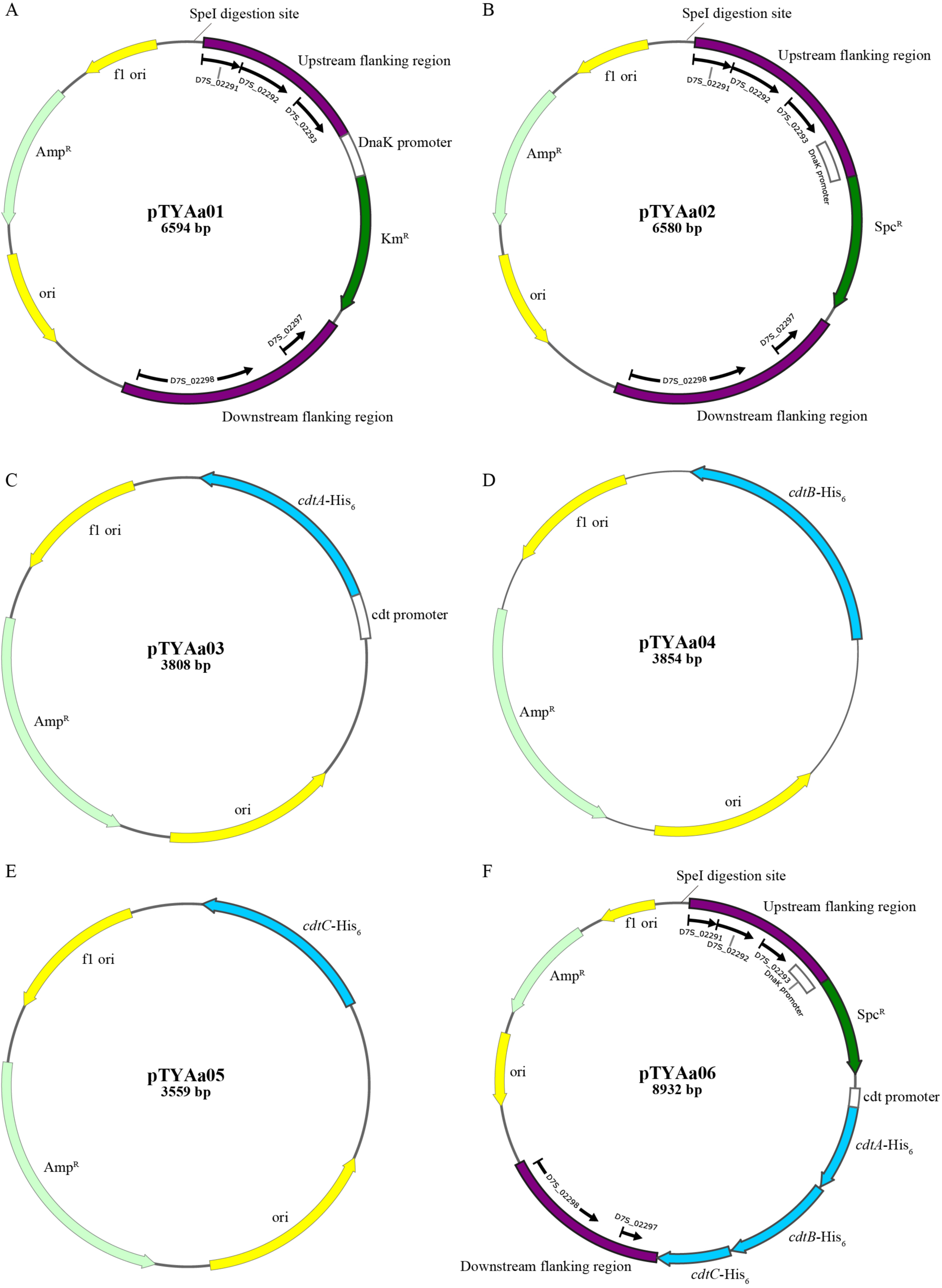
Plasmid construction maps for gene manipulation of *A. actinomycetemcomitans*. (Α) pTYAa01, designed to generate a *cdtABC* knockout with a kanamycin-resistance cassette via homologous recombination. **(B)** pTYAa02, designed to generate a *cdtABC* knockout with a spectinomycin-resistance cassette. **(C-E)** pTYAa03 (*cdtA*-His_6_), pTYAa04 (*cdtB*-His_6_), and pTYAa05 (*cdtC*-His_6_), cloned with 5’UTRs including the native promoter region of the *cdtABC* operon. **(F)** pTYAa06, designed to generate a *cdtABC* complement with a spectinomycin-resistance cassette via homologous recombination.

**Fig. S12.**
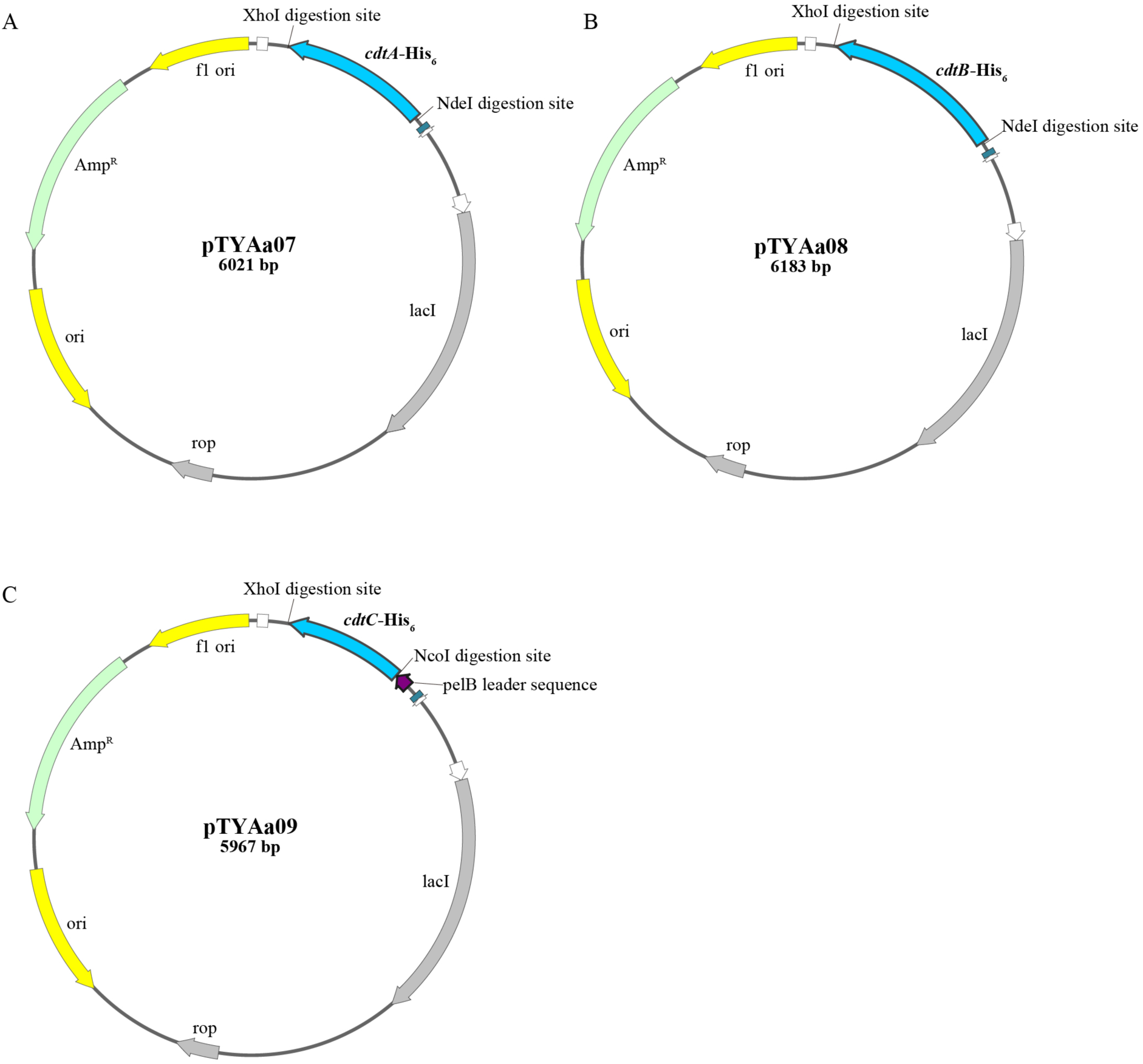
Construction of plasmids for the purification of recombinant CdtA, CdtB, and CdtC. **(A)** pTYAa07 for purification of His_6_-tagged CdtA. **(B)** pTYAa08 for purification of His_6_-tagged CdtB. **(C)** pTYAa09 for purification of His_6_-tagged CdtC.

**Table S1.**
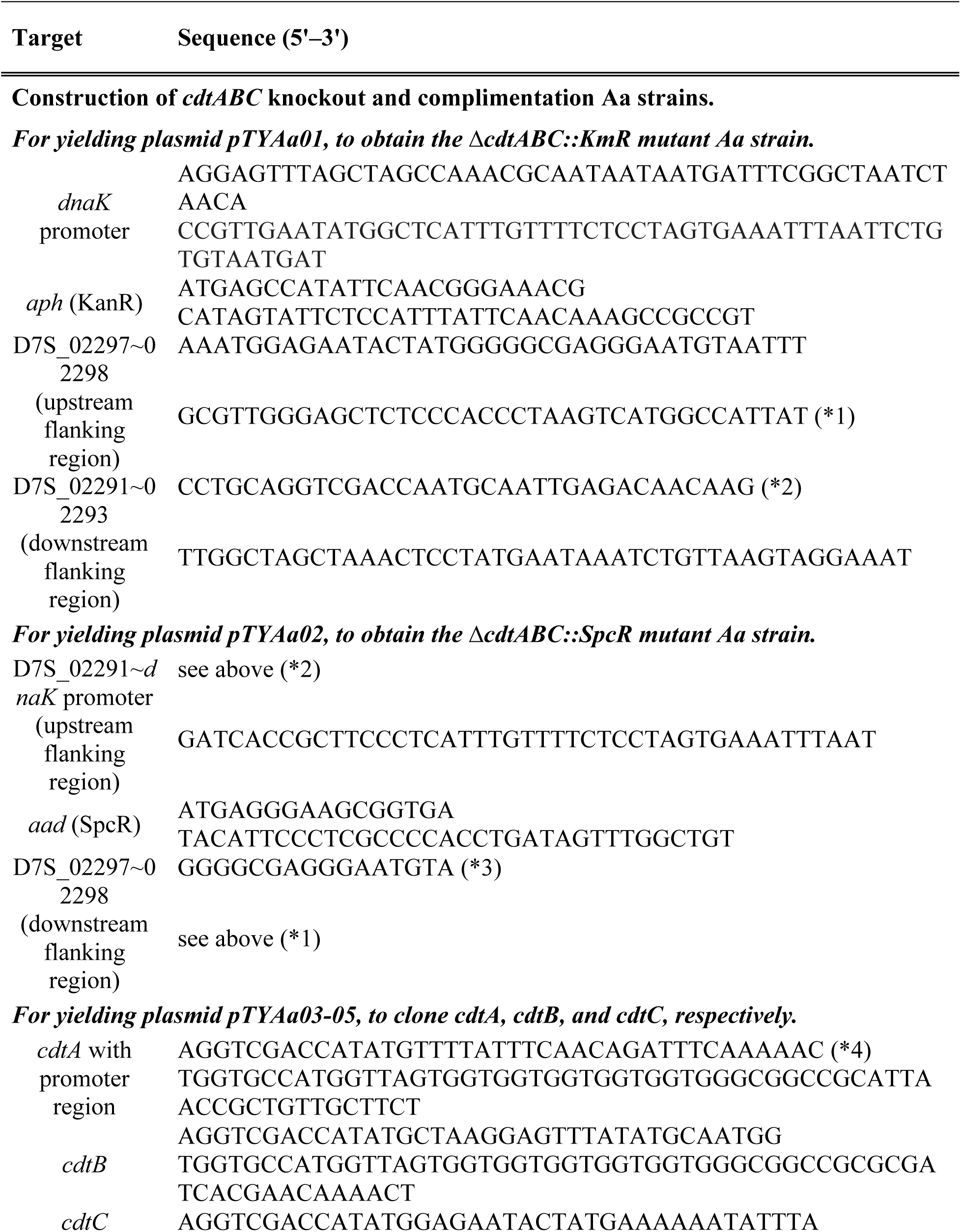
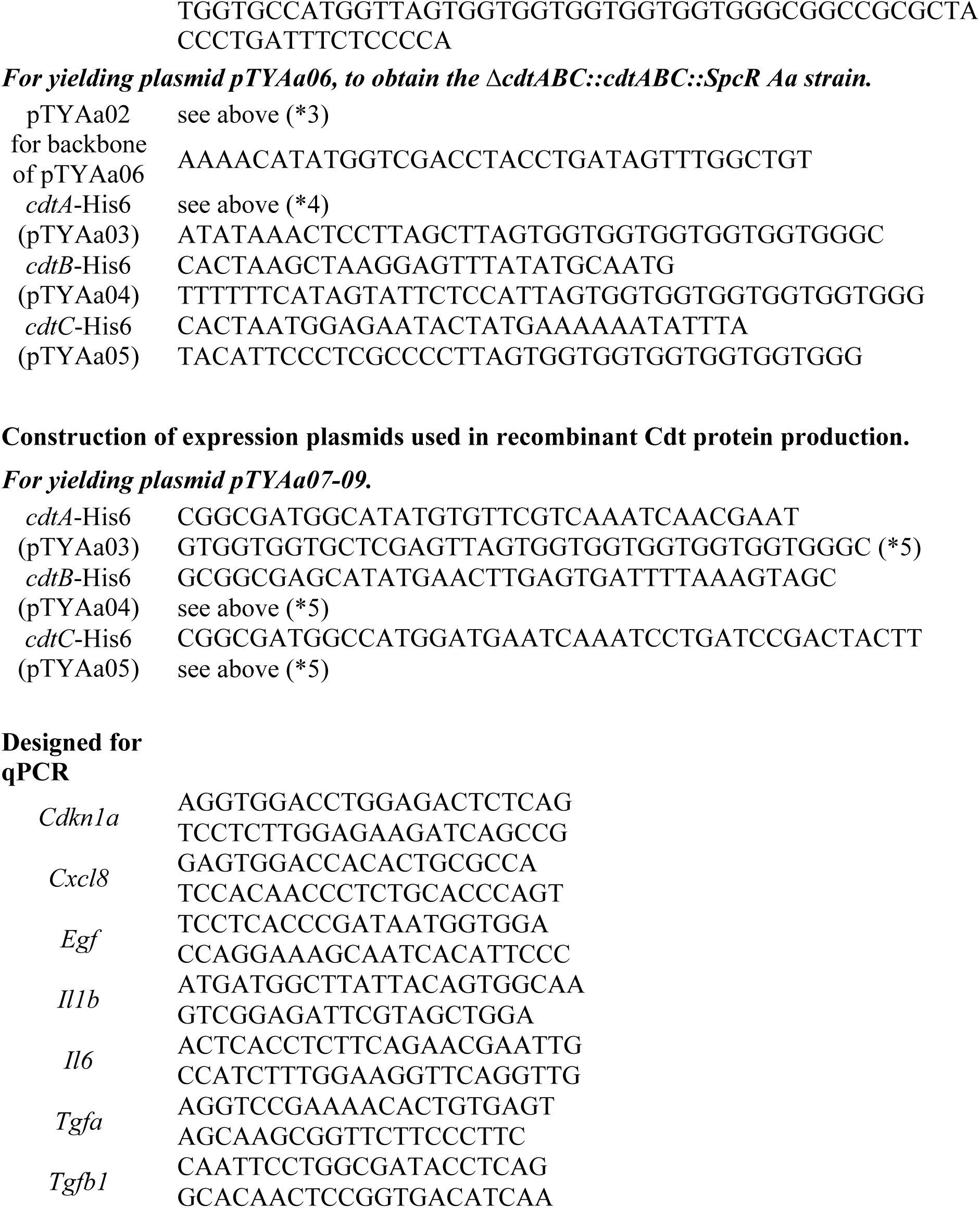
Oligonucleotides used in this study.

